# A novel platform of RNA 2′-O-methylation high-throughput and site-specific quantification tools revealed its broad distribution on mRNA

**DOI:** 10.1101/2020.03.27.011759

**Authors:** Yao Tang, Yifan Wu, Sainan Wang, Xiaolan Lu, Xiangwen Gu, Yong Li, Fan Yang, Ruilin Xu, Tao Wang, Zichen Jiao, Yan Wu, Liwei Liu, Jian-Qun Chen, Qiang Wang, Qihan Chen

## Abstract

Ribose 2′-O-methylation is involved in critical biological processes, but its biological functions and significance in mRNAs remain largely unknown due to the lack of accurate and efficient identification tools. To overcome this gap, we established NJU-seq (an accurate high-throughput single-base method) and Nm-VAQ (a site-specific quantification tool). We identified thousands of new Nm sites on mRNA of human and mouse cell lines, in which 68 of 84 selected sites were further validated to be 2′-O-methylated more than 1%. Unlike rRNA, the methylated ratios of most validated mRNA Nm sites were lower than 30%. In addition, mRNA 2′-O-methylation was dynamic-changing according to the circumstance, which was presented with MHV infection. Furthermore, Nm sites of lung surgery samples revealed commonness of lung cancer pathogenesis, providing potential new diagnostic markers.

## Introduction

More than 100 RNA modifications have been identified, constituting the epi-transcriptome code and affecting RNA functions (Motorin & Helm, 2011). Among these RNA modifications, 2′-O-methylation (otherwise known as Nm), whereby ribose is methylated at the 2′ hydroxyl position without limitation of the base identity, is a widespread RNA modification phenomenon, including multiple RNA types and species (Deryusheva et al., 2012; Jockel et al., 2012; Motorin & Helm, 2011; Tycowski et al., 1996; Zust et al., 2011). Furthermore, several sporadic published studies have demonstrated the importance of 2′-O-methylation in multiple biological processes as an essential signaling regulator. For example, Nm stabilizes the RNA structure by maintaining the ribose C3′-endo conformation, resisting nucleophilic attack, and avoiding 3′-5′ degradation triggered by uridylation (Ji & Chen, 2012; Kawai et al., 1992; Krogh et al., 2016). In addition, a single 2′-O-methylation modification as a molecular tag distinguishes between internal and external RNAs, generates abnormal alternative splicing, and inhibits translation (Ayadi et al., 2019; Daffis et al., 2010; Elliott et al., 2019; Ge et al., 2010; Lalonde et al., 2007; Ma et al., 2014; Zhao & Yu, 2008; Zust et al., 2011). However, despite its potential wide-ranging cellular activities, an in-depth exploration of 2′-O-methylation biological functions are still limited due to the lack of screening tools to detect Nm sites.

Three high-throughput Nm detection methods have been developed in the past few years: 2′OMe-seq (Incarnato et al., 2017), RiboMethSeq (Krogh et al., 2016), and Nm-Seq (Dai et al., 2017, 2018). These methods can reliably detect rRNA Nm modification for an abundant amount of rRNA and a high modification ratio. In addition, LC-MS and RTL-P were also used to validate the rRNA site-specific 2′-O-methylation status (Dong et al., 2012; Douthwaite & Kirpekar, 2007). However, the small amount, complex sequence context, and low methylation ratio of mRNAs lead to barriers to the practical application of those methods in mRNAs. To address this problem, we established an accurate high-throughput single-base Nm site screening method NJU-seq and a site-specific 2′-O-methylation quantification tool Nm-VAQ. With the help of these two methods, we efficiently revealed the broad distribution of Nm sites on mRNAs and observed that the 2′-O-methylation ratios of all validated Nm sites on mRNA were lower than 30%.

In the previous studies, rRNA was methylated by snoRNP complex, which contained FBL as methyltransferase, NOP56/58, and SNU13 as co-factor, partial complementary snoRNA with conserved C/D box as guide RNA (Yi et al., 2021). Although a particular Nm site on mRNA was reported to be methylated by the same mechanism (Elliott et al., 2019), how the rest of the Nm sites on mRNA methylated remained unknown. On the other hand, viruses such as HIV were reported to be 2′-O-methylated with human FTSJ3, which acts as a methyltransferase, in the host cell (Ringeard et al., 2019). However, further exploration of the whole mechanism relies on accurate Nm site information and methylation ratio evaluation.

## Results

### Establishment of NJU-seq

Based on the characteristic of 2′-O-methylation, our detection strategy was to take advantage of an Nm-sensitive RNase. Therefore, the newly established NJU-seq relied on helicase and hydrolase activities of RNase R from *Mycoplasma genitalium* (MgR) on ssRNA from the 3′ to 5′ direction, which produced the digested products end at every Nm+1 site (Lalonde et al., 2007). First, A synthesized 22 nt ssRNA substrate carrying one mixed 2′-O-methylation site NmNN (N=A/G/C/U) was incubated with MgR to evaluate its sensitivity to Nm sites (Figure 1A and Appendix figure 1A-B). As expected, the 18 nt ssRNA products accumulated with increasing MgR concentration, confirming the enzyme activity’s versatility. In addition, we also performed a reaction of MgR with the synthesized 40 nt ssRNA substrate specifically carrying only the Am site at position 28 to demonstrate the exact breakpoint of MgR activity, and the products were TA cloned and sequenced (Appendix figure 1C). Importantly, we discovered that all ssRNA products ended at the Am+1 position (20/20), supporting the detection of Nm sites by the accumulated 3′ end of MgR reaction products. To confirm that the digestion pause was stable in most contexts and may not be affected by certain secondary structures, we incubated an ssRNA with 2′-O-methylation at the 17th position among mixed ribonucleotides (N) from −10 to +10 position with MgR and the products were sequenced by NGS. As shown in Figure 1B, more than 93% of products ended at the Nm+1 position with almost equal ribonucleotides distribution from −10 to −1. In addition, we used multiple ssRNAs with an extended stem region as substrates. As expected, MgR digested all the ssRNA to the Nm+1 position successfully (Appendix figure 1E-F). In addition, an ssRNA substrate with m^6^A and m^5^C sites downstream of the Am site was incubated with MgR, demonstrating clear products ending at the Nm+1 site but nothing at other modification sites (Appendix figure 1D). All the above data indicated that MgR digestion was a great solution to create RNA fragments that ended at the Nm+1 position stably and precisely with no ribonucleotides, context, or secondary structure bias.

**Figure 1.**
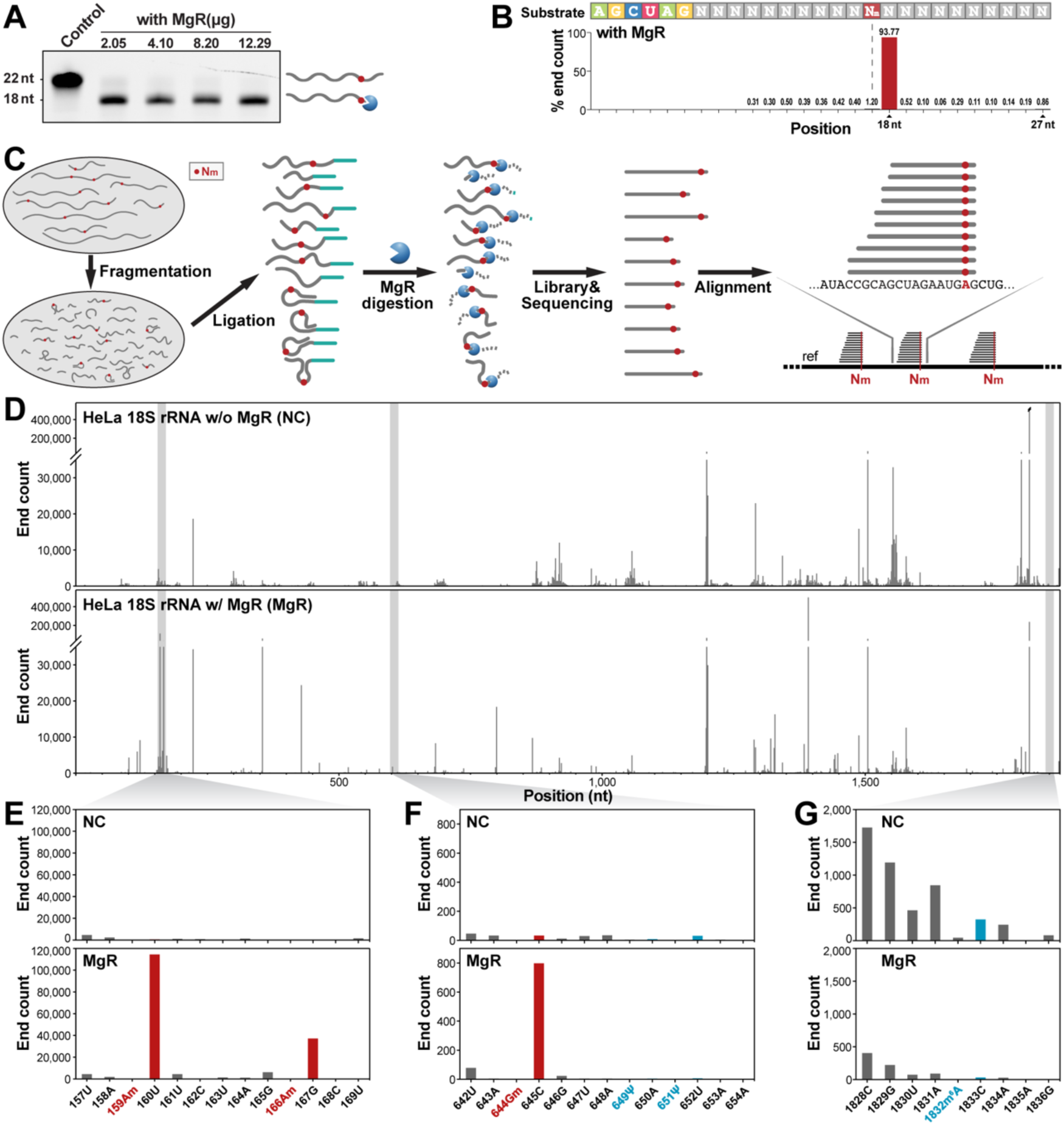
Establish a novel high-throughput method, NJU-seq, for RNA 2′-O-methylation. (A) *In vitro* ssRNA hydrolysis assay. The ssRNA substrate (5′-6-Fam-UAACCUAUGA AGNNNUNmNNC UC-3′, N represents the mix of A/G/C/U) was reacted with varying MgR amounts. The left electrophoresis showed digestion products of MgR, and a schematic was shown on the right. (B) High throughput sequencing of MgR hydrolyzed ssRNA products. The ssRNA substrate (5′-AGCUA GNNNN NNNNN NNmNNN NNNNN NN-3′) was incubated with MgR protein, followed by library construction and NGS. All reads shorter than 27 nt were counted, and the ratios were presented. (C) Schematic workflow of NJU-seq. 1) total RNA fragmentation; 2) 3′ tail ligation; 3) RNase R from Mycoplasma genitalium (MgR) digestion; 4) NGS library preparation and high-throughput sequencing; 5) alignment and analysis. (D) Plots showing read 3′ end counts on HeLa 18S rRNA with (MgR, bottom) and without (NC, top) MgR digestion. (E-G) Plots showing read 3′ end counts of regions 157-169, 642-654, and 1828-1836 on 18S rRNA, without and with MgR digestion. Sites marked with red were previously reported as 2′-O-methylated, and blue were previously reported as m^6^A or pseudouridine.

We combined MgR with NGS to develop a new high-throughput Nm screening method named NJU-seq (Nm-Judged-Universally sequencing). As detailed in the Methods, there were five main steps in NJU-seq (Figure 1C). A sample prepared with the same process without MgR treatment was used as a negative control (NC) group to filter the false-positive outcomes caused by RNA fragmentation bias. First, we tested NJU-seq on total RNA from HeLa cells and analyzed the rRNA results. The sequencing data were aligned to a reference sequence, and reads’ 3′ ends were counted (Figure 1D). Comparing rRNA alignments after MgR digestion to NC, 3′ end enrichments correspond to potential Nm sites. For example, between positions 157 to 169 in 18S rRNA, we found that the reads ending at positions 160 and 167 significantly accumulated, indicating two possible Am sites at positions 159 and 166 (Figure 1E). To further evaluate whether other RNA modifications on RNA would affect the activity of MgR, sites’ read distributions around all reported sites with eight other RNA epigenetic modifications were browsed, m^6^A, pseudouridine (ψ), ac^4^C, m^6^_2_A, m^1^A, m^7^G, m^5^C, and m^3^U (Appendix figure 2 and 3)(Taoka et al., 2018). Based on the results, none of those eight epigenetic modifications caused significant enrichment at 1nt downstream or surrounding other position comparing with NC, while Nm sites demonstrated opposite results as expected.

We scored each site with the difference of reads end 1 nt downstream divided by reads end just at the target site between MgR treatment and NC group to set a quantified standard (Appendix figure 5). The score was mainly correlated with the copy number of fragments with 2′-O-methylation modification at the Nm site (such as Nm-seq) but not the methylation ratio (such as RiboMethSeq). Candidate Nm sites were further judged by whether reads ending 1 nt downstream of the MgR treatment group were significantly more abundant than the NC group to exclude false-positive results by RNA fragmentation bias. The details of the scoring and filtering steps are described in Methods.

### Establish Nm-VAQ as a site-specific validation and quantification tool of Nm status

In the previous studies, RTL-P was widely used as a semi-quantification method that can validate sites with high 2′-O-methylation (Dong et al., 2012). However, RTL-P remained several disadvantages, such as hard to validate one site specifically or measure the methylation level accurately, results varying due to the number of targets, or demonstrating potential false positive or negative results (Dong et al., 2012). Here, we hope to develop a method to validate the 2′-O-methylation status of a specific site and acquire the methylation ratio. Hence, we established the Nm-VAQ (an RNase H-based Nm Validation and Absolute Quantification tool) with inspiration from previous relative studies (Yu et al., 1997). Briefly, Nm-VAQ was based on selective cleavage of RNase H and the guidance of a DNA-RNA hybrid probe to target the cleavage site (Figure 2A). With several tests of multiple designs of hybrid probes, we anchored the RNase H cleavage site to the target site with a probe containing 4 DNA and 13 RNA, as shown (Figure 2B and Appendix figure 4A-G). Combined with qPCR, Nm-VAQ quantified the mixed substrates with 2′-O-methylation gradients accurately with much lower methylation ratio (Figure 2C), while RTL-P could only detect Nm sites with modifications higher than 40% (Dong et al., 2012) and presented significantly different results under different conditions (Appendix figure 4H-I). In addition, the tests of Nm-VAQ with 50% 2′-O-methylated substrates at 6.25 fmol, 0.625 fmol, and 0.0625 fmol demonstrated highly consistent results, proving its evaluation capability in low-amount RNA studies (Figure 2D). To date, Nm-VAQ has allowed the study of Nm to the absolute quantification stage.

**Figure 2.**
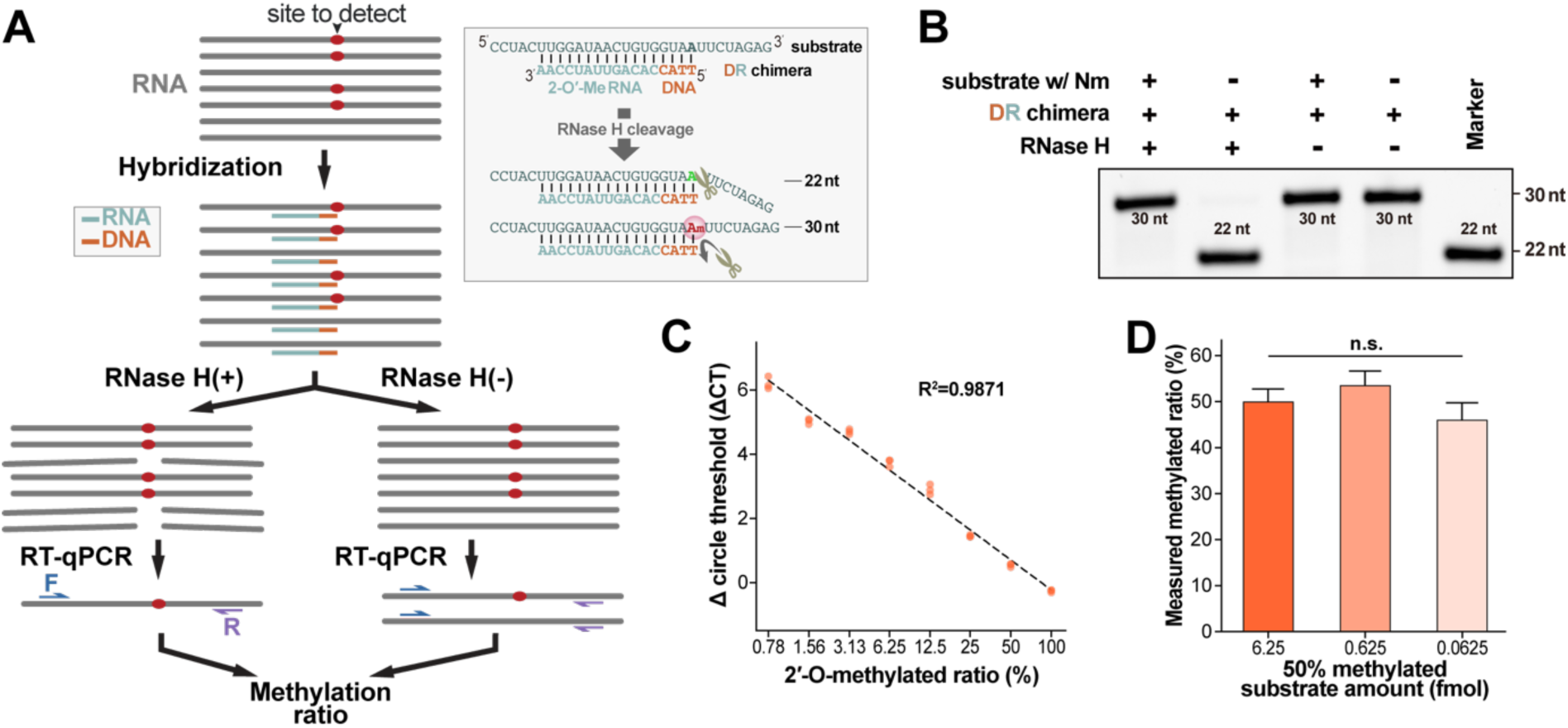
Establish a low-throughput Nm validation tool, Nm-VAQ. (A) Schematic workflow of Nm-VAQ. 1) the hybrid of RNA and chimera probes. In the substrate, the red site indicated the target site was 2′-O-methylated. In the chimera probe, the reddish-brown region showed the DNA, and the light green region showed the 2′-O-methylated RNA; 2) with/without RNase H cleavage; 3) RT-qPCR. The methylation ratio was from ΔCT (cycle threshold) of RNase H cleavage and control sample. The box on the right showed an example of RNase H cleavage directed by the chimera probe in Figure 2B. Hybrid schemes of RNA substrates (up sequences) and chimera probes (down sequences) were shown. (B) The RNase H reaction products were presented by electrophoresis. The number presented the length of FAM-labeled cleavage products. (C) Correlation between the 2′-O-methylated ratio of substrate and ΔCT (cycle threshold) for the Nm-VAQ assay. A 6.25*10^-2^ pmol substrate with and without Nm was mixed to obtain a known Nm ratio, and ΔCT was obtained from the RNase H treated and control samples. (D) Measurement of three different substrate concentrations with an Nm ratio of 50%. Nm ratio was calculated with ΔCT. n.s., not significant, by ordinary one-way ANOVA.

### Mapping of Nm sites of human cell lines

Although our main goal was to identify Nm sites on mRNA, we started from tRNA and rRNA to evaluate the results. As an example, NJU-seq identified Cm_32_ tRNA^Gln^ (UUG) turned out to be consistent with sites reported in previous studies (Marchand et al., 2017), which led to significant fragments accumulation end at 33th position (Appendix figure 6A). On the other hand, 83 Nm sites were identified on rRNA, including 81 Nm sites reported in the previous studies and 2 newly identified (Appendix figure 6B, Appendix table 1). Although NJU-Seq was not as sensitive as RiboMethSeq and LC-MS, it still identified approximate 3/4 of reported Nm sites on rRNA but with high specificity, which was reliable and promising to challenge the Nm identification on mRNA.

We next focused on the exploration of Nm sites on mRNAs from HeLa cell lines. With rRNA elimination, the proportion of mRNA in the substrate was substantially increased before MgR treatment. mRNA Nm sites shared with all three independent cell line samples were selected for the following analysis and statistics, as detailed in Methods. The same standard selected the potential Nm sites on mRNA as rRNA, which acquired 4074 in HeLa cell lines (Figure 3A and Appendix table 2).

**Figure 3.**
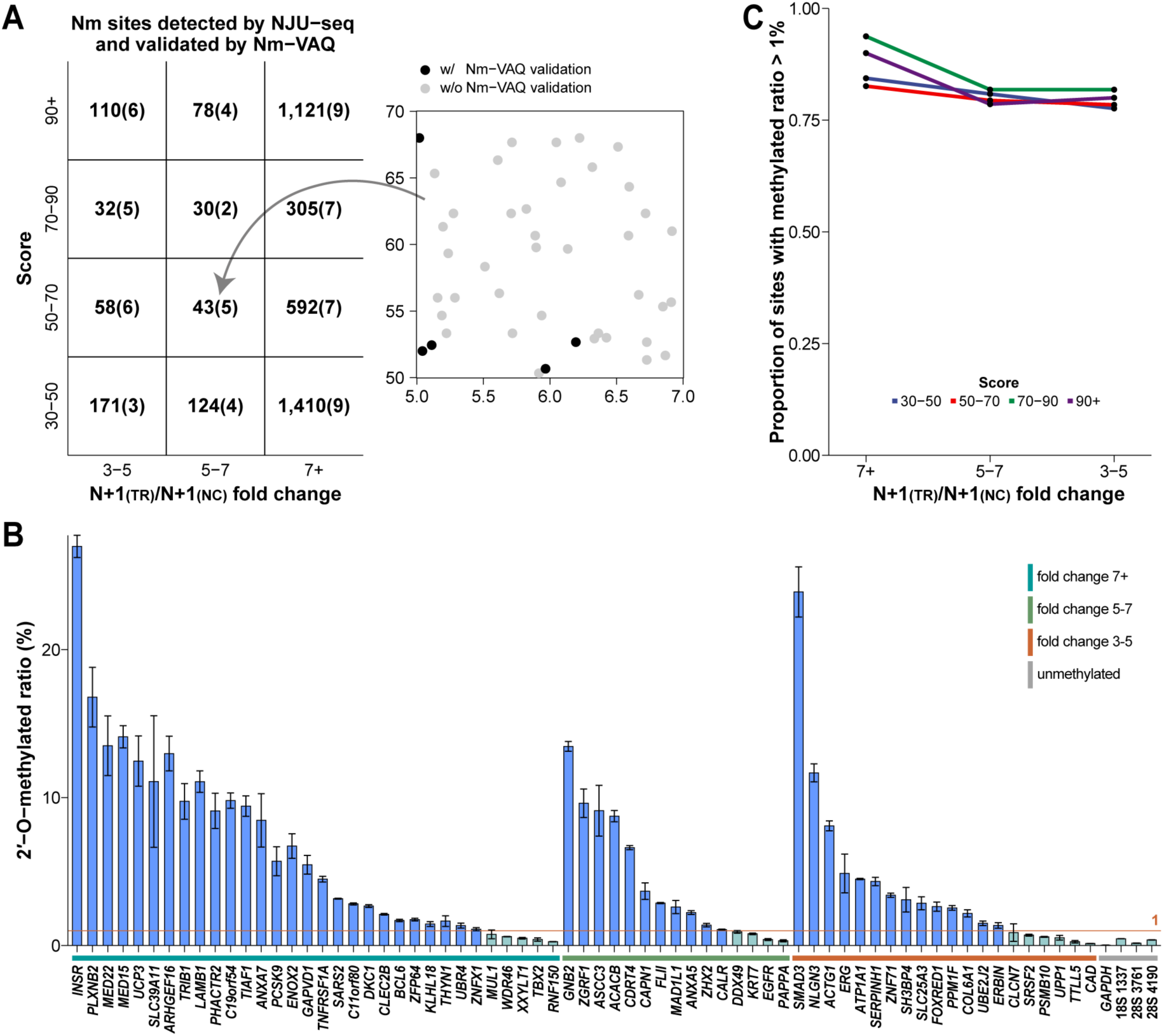
Detection and validation of HeLa mRNA Nm sites by NJU-seq and Nm-VAQ. (A) The distribution of HeLa mRNA Nm sites detected by NJU-seq and validated by Nm-VAQ. 4,074 mRNA Nm sites were screened in criteria the same as for rRNA. All Nm sites were divided into 12 regions based on score and foldchange. Each box represented the number of Nm sites detected by NJU-seq (without brackets) and by Nm-VAQ (in brackets), respectively (left). An example of the selection criteria for Nm-VAQ of Nm sites was shown on the right. As shown in the figure, Nm sites closest to score boundary and sites closest to fold change boundary were subjected to subsequent validation analysis by Nm-VAQ in each box, avoiding any artificial bias in the selection of validation candidates. The black dots indicated the selected sites for the following validation. (B) The methylation ratio in HeLa mRNA Nm sites detected by Nm-VAQ. The Nm sites are divided into three ranges of fold change (3-5, 5-7, 7+) and ranked from highest to lowest. The unmethylated sites on GADPH mRNA (12:6,537,190, G), and 28S 4190A, 3761C, 18S 1337C, which were reported as m6A, m5C, and ac4C in previous studies were selected as the negative control. The sites in blue indicated their 2′-O-methylation ratios were higher than 1%, while the cyan sites were lower methylated. The complete information of all selected sites can be found in Appendix table 6. Error bars described SEM for three technical replicates. (C) The proportion of sites with a methylated ratio higher than 1% in different fold change ranges and score ranges. Each of them was higher than 75%.

Before moving forward to further analysis and statistics of newly identified sites, we wanted to access the rationality of the standard we set and the approximate true positive rate of the 2′-O-methylated sites we selected. Generally, all 4,074 identified mRNA sites of HeLa cell lines were separated into 12 regions with the matrix of the score (30/50/70/90) and fold change (3/5/7) (Figure 3A). 67 sites were selected from the boundary regions of each region as the candidates to be further validated by Nm-VAQ (Figure 3A and Appendix table 2). As shown in Figure 3B, 52 of 67 sites demonstrated 2′-O-methylation ratios higher than 1%, while four negative control unmethylated or methylated by other modification sites demonstrated no positive results by Nm-VAQ. Meanwhile, the *in vitro* transcribed RNA fragments were used to evaluate the potential 2′-O-methylated status of relative sites on *PLXNB2* and *BCL6* by Nm-VAQ as a negative control, which did not produce any false positive signal (Appendix figure 7A). Besides, the *in vitro* transcriptome, which is similar amounts and composition of mRNA but with almost no modification, had been served as a negative control in RNA m^6^A modification detection tools to confirm the specificity of identified modification sites (Liu et al., 2022). Here, the mRNA site scores of *in vitro* transcriptome were significantly lower than those identified Nm sites of the template, indicating the high correlation of NJU-seq with Nm modification (Appendix figure 8). The methylation ratios of all selected sites were lower than 30%, much lower than sites in rRNA (Appendix figure 6C and Figure 3B). By calculating the proportion of sites with methylated ratios of each region and different standards, the specificity remained higher than 75% no matter which score threshold or fold change range we used (Figure 3C). Hence, NJU-Seq was confirmed to be a reliable method to screen potential Nm sites with pretty high accuracy.

We also selected 5 Nm sites on HeLa mRNA with high scores as targets for RTL-P validation; only three showed positive results compared with *GAPDH* as a negative control (Appendix figure 9A). Since the mRNA 2′-O-methylation status of most sites may be much lower than rRNA and RTL-P was limited by false positive, false negative, semiquantitative and non-single nucleotide resolution defects, Nm-VAQ was more appropriate in the following study than RTL-P.

We next applied NJU-seq on the other two cell lines, HEK293T and A549, and acquired 2,059/2,367 mRNA Nm sites. 1,044 mRNA Nm sites were shared among all three cell lines (Figure 4A). Three Nm sites shared by three cell lines were selected for further validation: *BCL6* (3, 187,721,452, Gm), *PLXNB2* (22, 50,290,219, Gm), and *TBX2* (17, 61,401,930, Cm). Nm-VAQ confirmed their methylation and evaluated 2′-O-methylation ratios as 0.4-18% (Figure 4B). All validated sites demonstrated clear 2′-O-methylation status but were much lower than Nm sites on rRNA. In addition, we performed GO enrichment analysis of genes with 3-cell-line-shared Nm sites and could see their extensive involvement in various biological processes (Appendix figure 10A).

To identify the overall distribution of 2′-O-methylation, we analyzed the sequence composition around the Nm site. We found that Nm modification occurred in a distinct pattern on different ribonucleotides but was similar among the three cell lines (Figure 4C). Although A on mRNA may act as a branch point in the intron part and regulate alternative splicing by its 2′-O-methylation status (Zhao & Yu, 2008), more 2′-O-methylations were modified on C and G than A and U. We found high GC content around the Nm sites in the three cell lines, especially G, which might indicate modification bias (Appendix figure 9B). When we further calculated Nm-1 and Nm+1 ribonucleotides with Nm together as a tri-ribonucleotide unit, the results showed an apparent uneven distribution with a clear enrichment of UGmU among all three cell lines (Figure 4C).

**Figure 4.**
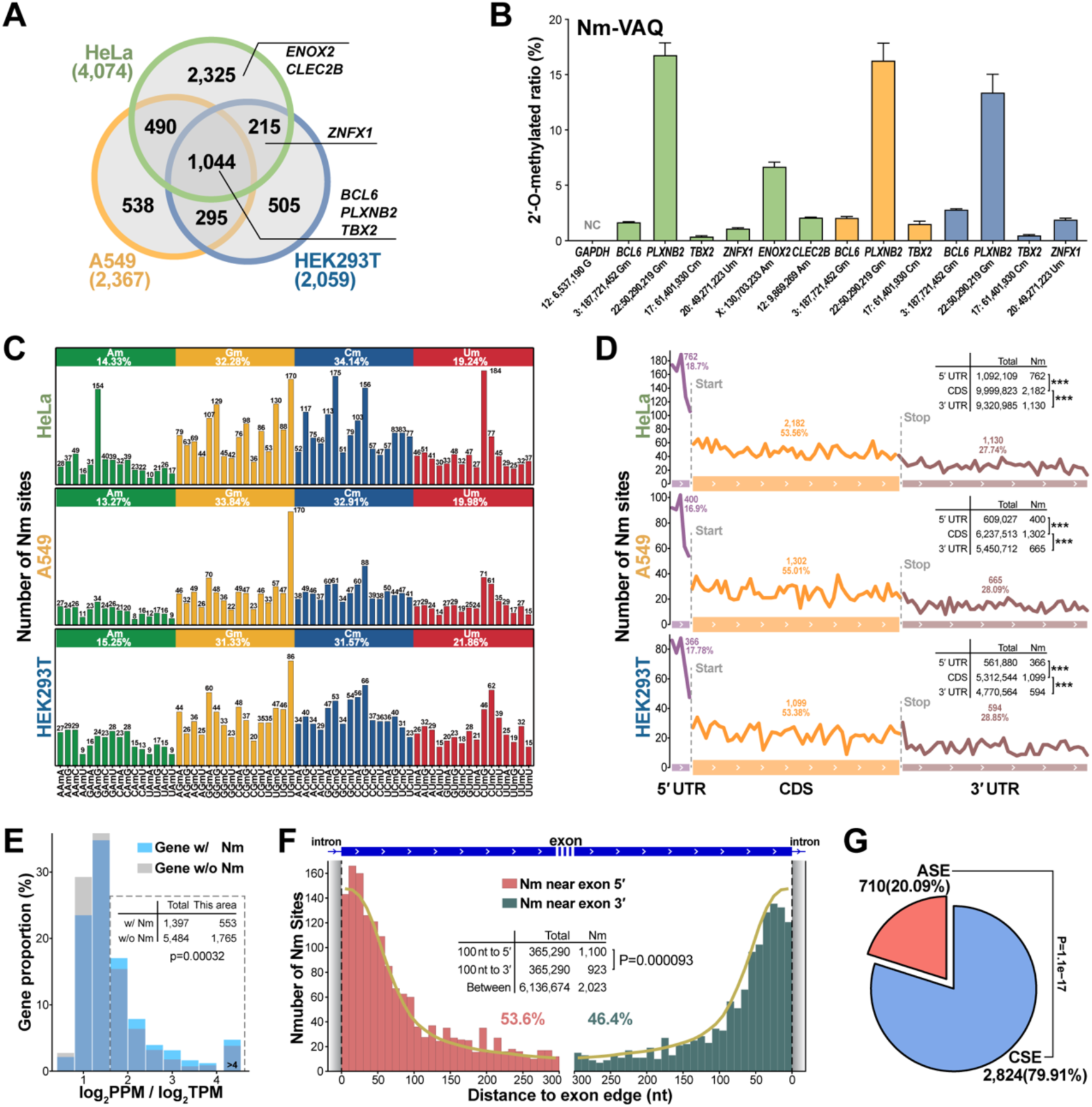
mRNA Nm sites in three human cell lines. (A) Overlap of mRNA Nm sites among the three cell lines by NJU-seq. (B) The methylation status in mRNA Nm sites detected by Nm-VAQ. The Nm sites of HeLa, HEK293T, and A549 cells were presented as green, blue, and orange sites, respectively. The HeLa Nm sites’ ratio data was the same as in Figure 4B. Error bars described SEM for three technical replicates. (C) Methylated nucleotide frequency distribution of HeLa mRNA. Nm-1, Nm, and Nm+1 sites were considered tri-ribonucleotide units. (D) Distribution of HeLa Nm sites on a length-normalized mRNA transcript (see Methods). ***P < 0.001 (Pearson’s chi-square test). (E) Translation efficiency of genes with and without Nm site(s) in HeLa. The ratio of protein abundance and RNA expression (log2PPM/log2TPM) represented translation efficiency. *P < 0.05 (Pearson’s Chi-square test). (F) Distribution of Nm sites near the exon 5′ (left) and exon 3′ (right) boundaries. All Nm sites were counted and divided into these two parts. The yellow dash represents the simulation of an even distribution. (G) Distribution of HeLa Nm sites in alternatively spliced exon (ASE) and constitutively spliced exon (CSE) regions. ***P < 0.001 (Pearson’s chi-square test).

### 2′-O-methylation is involved in mRNA alternative splicing events and translation

To identify the potential functions of 2′-O-methylation modifications, we analyzed Nm′s distribution patterns in different functional mRNA regions. By exploring the 2′-O-methylation distribution pattern on mRNA transcripts, we found that in the three cell lines, Nm numbers were CDS>3′UTR>5′UTR, which demonstrated the highest density in the 5′UTR considering the length (Figure 4D). To further investigate the Nm site distribution, we focused on the start and stop codons (Appendix figure 9C-D). 3,479 HeLa Nm sites on mRNAs that could cover −50 to +500 nt around the start codon were selected to count the distribution in 50nt-wide windows. Similarly, 3,074 HeLa Nm sites were selected to evaluate the stop codon’s −500 to +500 nt regions. Interestingly, both areas demonstrated apparent downward trends of Nm sites from 5′ to 3′ direction. Such a pattern was also seen in the other two cell lines.

Such results implied that there were relatively stable patterns of Nm distribution in the functional regions of mRNA. Next, we compared the translation efficiency of genes with and without 2′-O-methylation (Figure 4E). The translation efficiency was reflected by protein abundance/gene expression. Significantly more genes with Nm site(s) showed higher translational efficiency, which strongly suggests a correlation between Nm sites and translation, confirming the vital role of 2′-O-methylation in translational regulation. Previous studies elucidated that pre-mRNA splicing of *ACT1* was affected by Am modification, so we explored whether Nm sites were extensively involved in mRNA splicing (Zhao & Yu, 2008).

Interestingly, there were more Nm sites closer to the 5′ exon boundary than those closer to the 3′ exon boundary, significant in the 0-100 window from the boundary (Figure 4F and Appendix figure 9E,G). Such discoveries positively indicated the involvement of 2′-O-methylation in mRNA splicing. To further demonstrate the effects of splicing, we assigned mRNA Nm sites as constitutively spliced exons (CSEs) or alternatively spliced exons (ASEs) (Ge et al., 2010; Ryan et al., 2012). The total length of ASE and CSE with modifications was counted, and the density of Nm sites was obtained. We found that more Nm sites were significantly located in the CSE than in the ASE when considering their different lengths (Figure 4G and Appendix figure 9F,H). Such results suggested the importance of 2′-O-methylation in the occurrence and regulation of splicing during mRNA maturation.

### 2′-O-methylation dynamic response to virus infection

As Nm sites were associated with gene functions, it was interesting to explore whether mRNA 2′-O-methylation participates in gene regulation in response to external stimuli. Therefore, we designed virus infection experiments to observe the dynamic change in Nm sites. We first selected the mouse neuroblastoma (Neuro-2a) cell line as a target to explore mRNA 2′-O-methylation changes before and after mouse hepatitis virus (MHV) A59 infection as a model of viral infection response (Figure 5A). Based on the same standard as the previous parts, 2,272 and 2,549 Nm sites were identified from Neuro-2a cell lines with or without MHV infection, among which 496 Nm sites specifically appeared in the infection group (Figure 5B and Appendix table 3).

**Figure 5.**
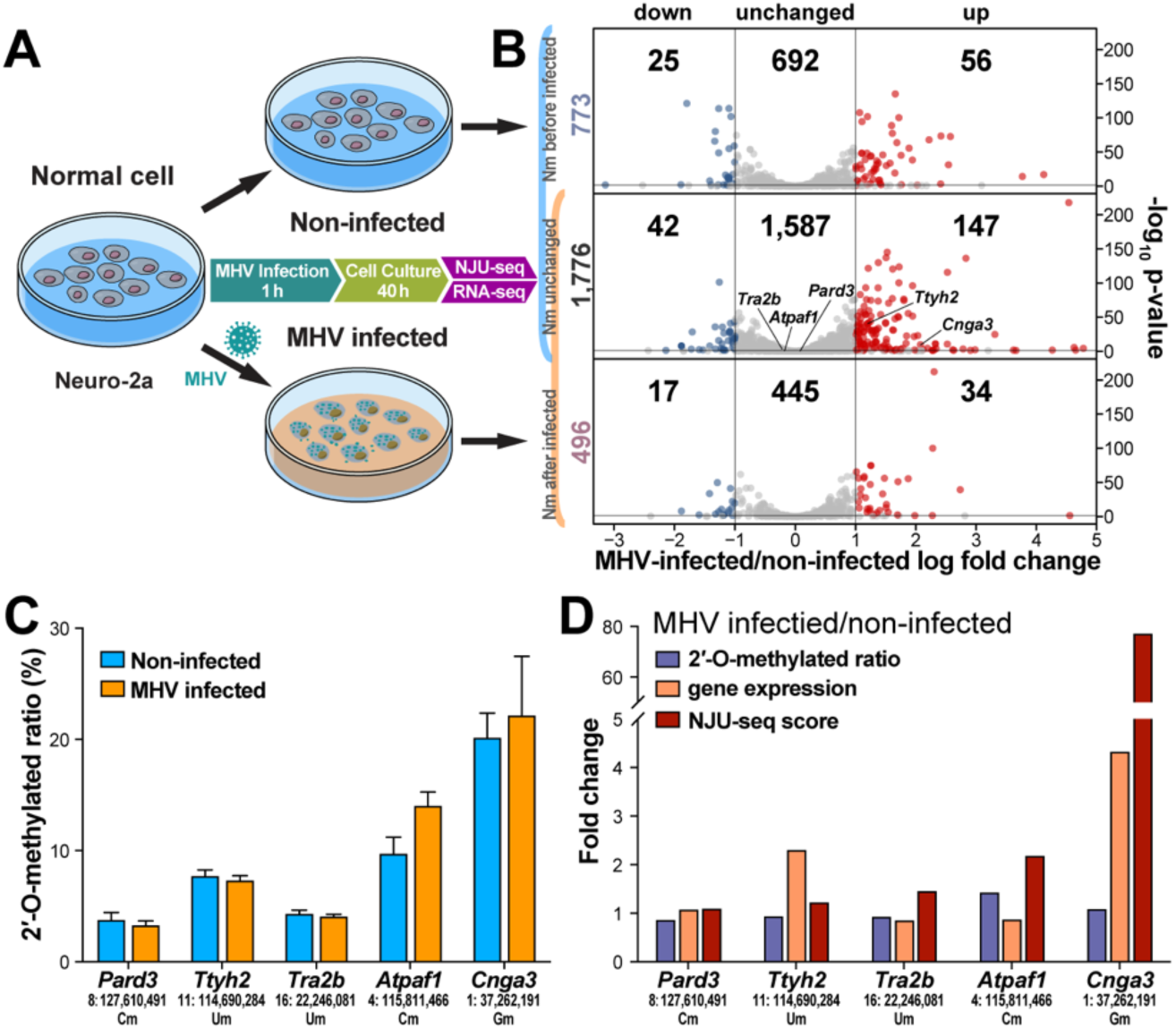
Dynamic 2′-O-methylation response to virus infection. (A) mRNA Nm sites changed in the Neuro-2a cell line after MHV infection. The flow of MHV infection was shown. Each of three MHV-infected and non-infected samples was performed with NJU-seq and RNA-seq. (B) 773 non-infected-specific, 1,776 shared, and 496 MHV-infected-specific Nm sites were detected as shown left. Genes containing these Nm sites were identified as down-regulated, unchanged, and up-regulated according to RNA-seq. (C) The 2′-O-methylated ratio of shared Nm sites detected by Nm-VAQ. Error bars described SEM for three biological replicates. (D) Fold changes of the shared Nm sites’ 2′-O-methylated ratio, NJU-seq score, and Nm-site-located genes’ expression after MHV infection.

Together with the results of RNA-Seq, all sites and the relative mRNA expression were divided into nine groups with MHV-infection specific/shared/non-infection specific and expression down-regulated/unchanged/up-regulated as shown (Figure 5B and Appendix figure 10B-C). Five sites were selected to evaluate its 2′-O-methylation status and expression change further. Among them, the methylation ratios of *Pard3* (8:127,610,491, Cm), *Ttyh2* (11:114,690,284, Um), *Tra2b* (16:22,246,081, Um) and *Cnga3* (1:37262191, Gm) showed no obvious change, while the methylation ratio of *Atpaf1* (4:115,811,466, Cm) increased from 9.8% to 14.1% (Figure 5C-D).

Based on the principle of NJU-Seq, the results were affected by both methylation status and expression change. For example, the expression level of *Atpaf1* remained no apparent change, while the methylation ratio increased by about 4%, which led to a significant increase in the NJU-seq score. On the other hand, the methylated ratio of *Cnga3* remained at approximately 20%, but the score by NJU-seq increased dramatically due to 4 times more up-regulated mRNA expression (Figure 5D).

### Nm demonstrated significant different patterns in normal and lung cancer tissues

Since the viral infection was a transient change, we further explored the dynamics of 2′-O-methylation in chronic diseases. Here, we used eight lung cancer clinical samples (6 lung adenocarcinoma (LUAD) and 2 lung squamous cell carcinoma (LUSC)) as a chronic disease response model. NJU-seq was performed to study Nm site differences in lung cancer tissues compared to distant normal lung tissues. As indicated, normal tissue-specific Nm sites were more often present in only 1 to 3 samples, while cancer samples had more shared Nm sites, implying the commonness of lung cancer pathogenesis (Figure 6A and Appendix figure 10D). Such cancerous-specific mRNA Nm sites are ideal candidates as diagnostic markers and are essential in future cancer studies (Figure 6B, Appendix figure 11D-I, and Appendix table 4). For instance, *CDH22* encodes the transmembrane protein cadherin-like protein 22, crucial in metastasis or the spread of cancer (Aydin et al., 2019; Piche et al., 2011). Cm (20:46,174,611) located in *CDH22* CDS was detected in all eight cancer samples but not distant normal samples with no correlation to mRNA expression differences.

**Figure 6.**
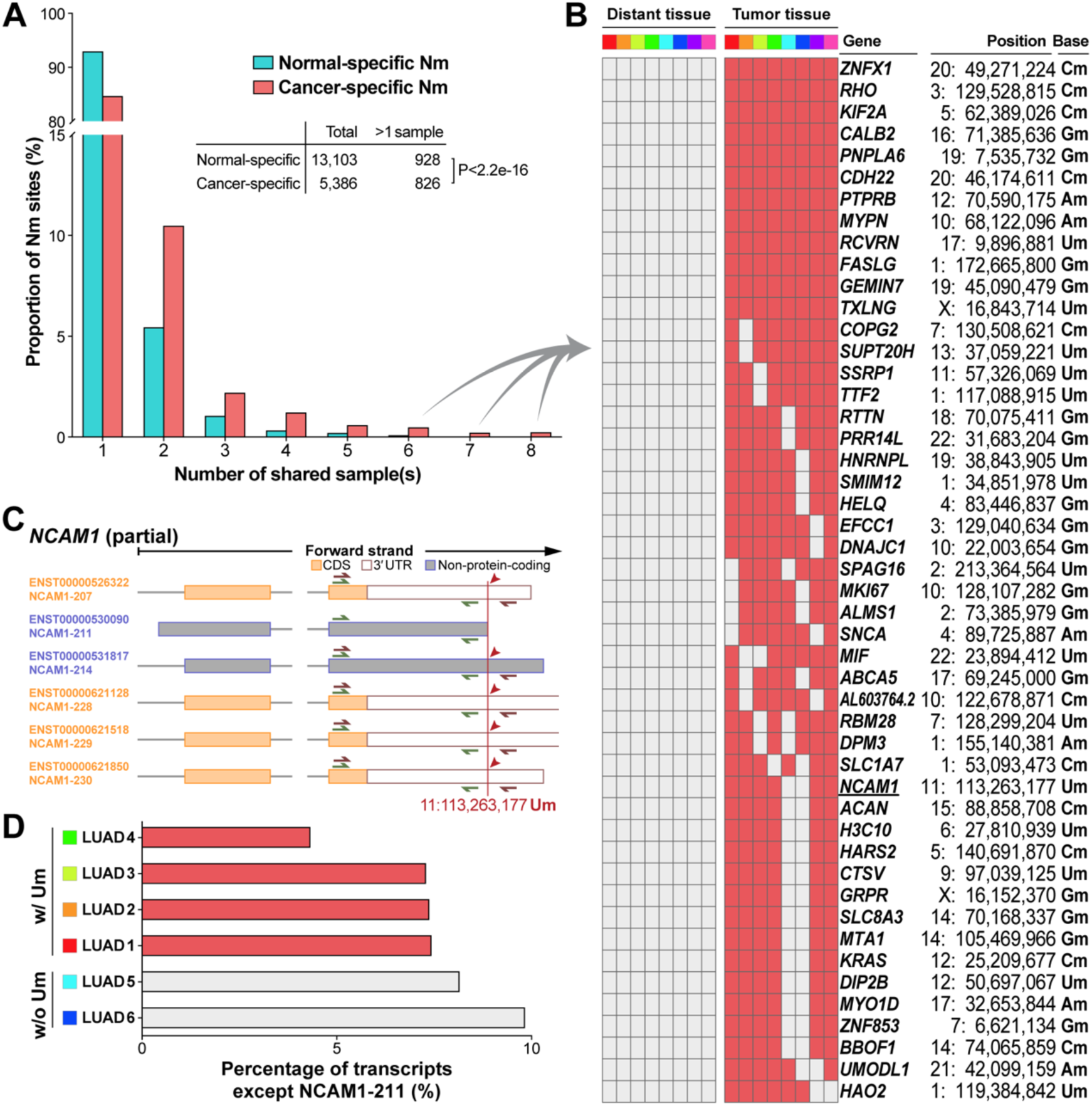
Significant different Nm patterns in normal and lung cancer tissues. (A) Sharing of specific Nm sites detected in lung cancer or normal tissue samples was shown. Eight pairs of tumor tissues and distant normal tissues from different lung cancer patients were sequenced and detected. Cancer-specific Nm sites occurred in at least one cancer tissue but were not detected in any normal tissue. The opposite is true for normal-specific Nm sites. The numbers of sites specific in over one cancer or normal tissue were subjected to the chi-square test. (B) Specific mRNA Nm sites among over five lung cancer samples are listed. (C) Um site (11:113,263,177) on *NCAM1* was located at the 3′ end of the transcript NCAM1-211. The green or red arrows indicated the primers for six transcripts (NCAM1-207, 211, 214, 228, 229, and 230) or five transcripts (without NCAM1-211) RT-qPCR amplification. (D) Percentage of transcripts except NCAM1-211. The proportion of the above five transcripts to six transcripts covering the Nm site was shown. Pastel red means Nm-detected, while gray means undetected.

Further analysis of cancer-specific Nm sites revealed that Nm sites might regulate the molecular function of cancer gene mRNAs in multiple ways. As mentioned above, Nm is closely related to splicing, so we focused on regulating the splicing complex. Interestingly, several Nm sites located upstream of the alternative splicing site can be found, regulating alternative splicing products. For instance, the *NCAM1* gene is involved in cell growth, differentiation, and migration (Sasca et al., 2019). Um (11:113,263,177) on *NCAM1* mRNA was detected in 4 LUAD and 2 LUSC samples located on the 3′ boundary of transcript NCAM1-211 (Figure 6C). We examined all six transcripts covering this site (NCAM1-207, 211, 214, 228, 229, 230) to verify the association between the Um site and splicing. Among 6 LUAD samples, the expression ratio of 5 longer transcripts was lower in 4 LUAD samples with Nm than in 2 samples without Nm, suggesting that 2′-O-methylation modification was associated with alternative splicing to regulate the mRNA products (Figure 6D and Appendix figure 11J-M).

## Discussion

The 2′-O-methylation modification of RNA ribose affects diverse biological processes. Thus, high-throughput identification for comprehensive and accurate profiling of Nm sites is vital for further research on cellular regulation and functions of 2′-O-methylation. Three high-throughput Nm detection methods have been developed in the past few years: 2′OMe-seq (Incarnato et al., 2017), RiboMethSeq (Birkedal et al., 2015; Erales et al., 2017; Krogh et al., 2016), and Nm-Seq (Dai et al., 2017, 2018) (Table 1). Based on this principle, 2′OMe-seq and RiboMethSeq could identify Nm sites with abundant amounts and high methylation ratios with very good sensitivity such as most Nm sites on rRNA, but various and inconsistent results on Nm sites with lower methylation ratio. For example, 18S 354U was ∼20% 2′-O-methylated, which was identified as methylated in 2016 by 2′OMe-seq, but the opposite by RiboMethSeq (Incarnato et al., 2017; Krogh et al., 2016). Such challenge was also seen on 18S 1440 Cm, which was not reported until 2020 (Krogh et al., 2020). Based on our results, the methylation ratio of almost all tested mRNA Nm sites was lower than 30%, which indicated that most Nm sites on mRNA were unable to be detected by 2′OMe-seq and RiboMethSeq. On the other hand, the Nm-seq method demonstrated improved mRNA Nm site detection direct targeting to mRNA molecules with 2′-O-methylation (Dai et al., 2017, 2018), similar to our new method NJU-seq. However, two main problems remained and were solved by NJU-seq: 1) Nm-seq based on multiple cycles of OED+RNA purification to enrich the Nm signal, which resulted in a massive loss of RNA content and took at least 16 hours. For NJU-seq, enrichment of the Nm+1 site can be accomplished with only 30 min MgR digestion for only one reaction with much less RNA sample. 2) Ligation bias of fragments. After Nm-seq treatment, the 3′ end of the RNA was 2′-O-methylated, resulting in a much lower ligation efficiency than fragments without oxidation process (Saikia et al., 2006). Thus, some Nm sites were insufficient to obtain more sequencing reads than surrounding non-modified sites. The RNA fragment of NJU-seq ended at the Nm+1 site without 2′-O-methylation, which has no bias in the ligation step. Those shortages may limit the use of Nm-seq in subsequent studies.

**Table 1.**
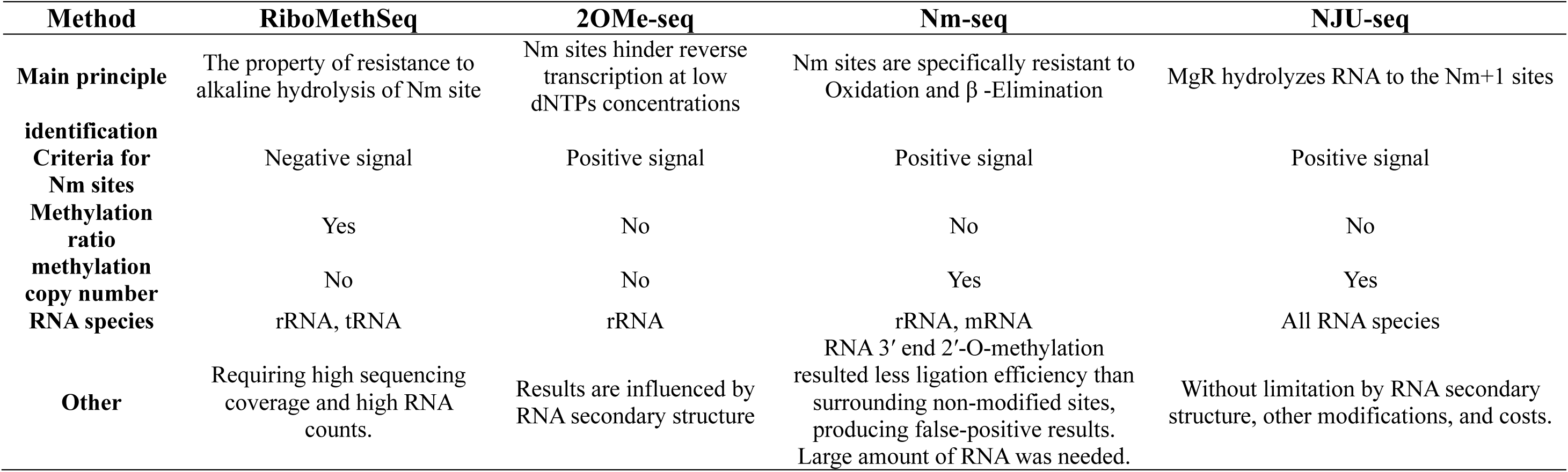
Comparison of 2′OMe-seq, RiboMethSeq, Nm-seq, and NJU-seq

To observe the specificity and sensitivity of NJU-seq, we knocked known rRNA 2′-O-methyltransferase fibrillarin (*FBL*) by siRNA in the HeLa cell line as reported in the previous study (Elliott et al., 2019; Erales et al., 2017). Although *FBL* knockout was proved to be lethal, siRNA could lower the expression of *FBL*, which led to a small decrease of rRNA 2′-O-methylation (Erales et al., 2017). As expected, *FBL* mRNA expression was significantly decreased that confirmed by RT-qPCR (Appendix figure 12A). After normalization to total rRNA-aligned reads, fragments ending 1 nt downstream reported Nm sites in the *FBL*-KD group were significantly fewer than those in the untreated group (Appendix figure 12B-C). Such results supported that NJU-seq can reflect the copy number of fragments with 2′-O-methylation. Besides, the methylation levels of several selected sites previously reported as Nm sites between the HeLa negative control and *FBL*-KD groups decreased by 8-20%, as quantified by Nm-VAQ (Appendix figure 12D).

The site-specific Nm validation and quantification method Nm-VAQ was another new tool established in this study to fulfill the Nm quantification gap in the study of 2′-O-Methylation. Several previously developed low-throughput Nm detection methods, including LC-MS, RTL-P, and DNA polymerase, have various defects (Aschenbrenner & Marx, 2016; Dong et al., 2012; Douthwaite & Kirpekar, 2007). LC-MS is labor-intensive and difficult for mRNA Nm detection due to the requirement for a large number of RNA molecules (Douthwaite & Kirpekar, 2007). Both RTL-P and DNA polymerase relied on Nm blocking on reverse transcription, which can be called RT-based methods (Aschenbrenner & Marx, 2016; Dong et al., 2012). Any Nm site between the amplification products will generate a methylation signal, and thus, these two detections are non-site-specific methods. Meanwhile, although both RTL-P and DNA polymerase methods could acquire linear results correlated with the methylation ratio with synthetic RNA, the result varied with different amounts of target RNA. As we observed in the evaluation of 18S 1391Cm, the results were significantly different in three RNA amounts, making it difficult to accurately evaluate the 2′-O-methylation ratio of the actual target (Appendix figure 4I). In addition, the original study of RTL-P mentioned false positive and negative results, which may be caused by RNA secondary structures that can occur on several rRNA sites (Dong et al., 2012). Nm-VAQ demonstrated apparent advantages compared to those methods, as listed in Table 2. Nm-VAQ anchored the cleavage position to the target site directed by the chimera and discriminated 2′-O-methylated RNA from unmethylated RNA molecules by RNase H. This method simultaneously acquired the absolute amount of accurate 2′-O-methylation of the target site. In addition, Nm-VAQ showed its capability to consistently evaluate targets with low amounts or low methylation ratios, demonstrating its capability to study the status of specific Nm sites and their dynamic changes.

**Table 2.** Comparisons of LC-MS, RTL-P, DNA polymerase, and Nm-VAQ methods

| Method | LC-MS | RTL-P | DNA polymerase | Nm-VAQ |
| --- | --- | --- | --- | --- |
| Main principle | Purification of RNA fragments containing modification sites and specific enzymatic hydrolysis to separate and analyze modification signal | 2'-O-methylation induced reverse transcription stop at low dNTP concentrations | Am or Cm in 1nt downstream of the primer hindered the reverse transcription reaction | 2'-O-methylation inhibited the cleavage of RNase H |
| Region/site-specific | Yes | No | No | Yes |
| Effective target | Am/Gm/Cm/Um | Am/Gm/Cm/Um | Am /Cm | Am/Gm/Cm/Um |
| Methylation ratio | Yes | No | Yes | Yes |
| absolute quantification of methylation copy number | No | No | No | Yes |
| Potential false result | No | Yes | Unknown | No |
| Other | Large amounts of high purity RNA, which is very laborious, are not suitable for detecting modifications of low abundance RNA | The methylation ratio varied significantly with the different RNA expression |  | Precise quantification of Nm ratio and methylation copy numbers at unknown RNA concentrations |

In the data process, we set two parameters score and fold change as 30+ and 3+ with a higher standard to maintain a high true positive rate of the screened Nm sites, which was later confirmed to be higher than 75% by Nm-VAQ. More restricted parameter would theoretically result in higher true positive rate, which means both parameters could be adjusted due to the purpose of the study. Although some screened sites were proved to be methylated with lower than 1%, the amount of methylated fragments was not negligible considering their high expression level (Appendix figure 7B), which might play essential roles in cell metabolism. Furthermore, NJU-seq achieved high consistency in mRNA duplicate samples (Appendix table 2 and Appendix table 3). These results indicated the stable broad-spectrum applicability of NJU-seq in detecting Nm modification in various RNA molecules with high complexity and low content.

In this study, we identified approximate 3/4 previous reported as well as two new Nm sites on rRNA of HeLa cell line. 18S 354U and 1440C were further validated to be 2′-O-methylated with ratio lower than 50%, which may be hard to detect consistently by RiboMethSeq and LC-MS. By screening the snoRNA database, 18S 1320G and 1784G could be targeted by snoRNA117 and snoRNA116-4, which need further validation in the future study. On mRNA, thousands of new Nm sites were uncovered on mRNAs from humans and mice, installing multiple areas (UTR/CDS, CSE/ASE), revealing its broad distribution. Unlike Nm sites on rRNA, all tested Nm sites on mRNA were 2′-O-methylated at less than 30%, which was hard to answer whether such low methylation ratios could regulate biological processes widely. However, based on previous studies, m^6^A were proved to be modified with a median modification ratio of ∼40% that was enough to regulate the function of mRNA (Liu et al., 2022; Liu et al., 2013). On the other hand, NJU-seq and Nm-VAQ studied a group of cells from cell lines or tissues, it was unclear whether all cells demonstrated such low modification or some cells were highly methylated while others were not, which required other methods to explore further. A similar distribution pattern of 2′-O-methylation was shared in different cell lines. The non-random distribution of Nm sites and the occurrence of 2′-O-methylation were tightly regulated in critical functional regions, suggesting that Nm might be an essential regulatory means. Thus, 2′-O-methylation modifications played an essential role in mRNA metabolism. In contrast to m^1^A with enrichment in the start codon or m^6^A with accumulation around the stop codon (Dominissini et al., 2016; Li et al., 2016; Meyer et al., 2012), the Nm distribution pattern was more complicated than these two RNA modifications and may possess broader biological functions. Consistent with Nm-Seq results, we did not discover any conserved motif for mRNA 2′-O-methylation (Dai et al., 2017, 2018). Such results implied mRNA 2′-O-methylation went through a different pathway than sequence-specific 2′-O-methylation on rRNA by FBL/snoRNA or structure-specific 2′-O-methylation on tRNA/miRNA/piRNA by FTSJ and HENMT1 (Liang et al., 2020; Marchand et al., 2017). Based on the novel platform demonstrated in this study, the distribution and quantification of Nm sites were acquired broader and easier, which provide an important basis for the follow-up mechanism exploration.

Based on the limited results, it was interesting that 2′-O-methylation might affect the interaction between mRNA molecules and proteins by affecting binding. According to previous studies, 2′-O-methylation could affect RNA-protein interactions by changing the binding surface and space (Jockel et al., 2012; Zust et al., 2011). Since AGO-miRNA-mRNA regulation is one of the most well-known mechanisms, we further analyzed the potential interaction between the RNA-induced silencing complex (RISC) and Nm sites in an infected Neuro-2a cell line. In the MHV-infected Neuro-2a cells, Gm (15:5,097,989) of *Card6* mRNA was located upstream of the cleavage site precisely in the target area of mmu-miR-3073a-3p (Appendix figure 11A-C and Appendix figure 13A,C), which may inhibit RISC cleavage, as reported in a previous study (Martinez & Tuschl, 2004). *Card6* belongs to the CARD gene family, is induced by interferon in multiple immune cells, regulates the immune response, and plays a vital role in inflammation (Li et al., 2014). In addition, we mimicked the 2′-O-methylation on the target mRNA of RISC based on the RISC-mRNA crystal structure (PDB 6MDZ) and the distances between the relative amino acid and modified ribonucleotides of positions 5, 12, and 20 on mRNA were too short, which may interfere with RISC binding (Appendix figure 13B). In addition, other biological processes relying on RNase H-like RNA endoribonuclease activity would all be inhibited by 2′-O-methylation at particular sites, such as Prp8 in alternative splicing, Dicer in miRNA maturation, and RNase L in RNA cleavage of the immune response (Burke et al., 2019; Fica et al., 2017; Ha & Kim, 2014).

With the help of NJU-seq and Nm-VAQ, further studies about the function, modification/de-modification/regulation of 2′-O-methylation could be extended. On the other hand, 2′-O-methylation demonstrated a critical function in environmental response and great potential in disease study, diagnosis, and drug discovery.

## Materials and METHODS

### Cloning, expression, and purification

The RNase R sequence from *Mycoplasma genitalium* (accession number: WP_009885662.1) was synthesized by GenScript Biotech Co. and inserted into the pET28a vector (+) with digestion and ligation at the NdeI and BamHI sites, followed by confirmation by Sanger sequencing. The plasmid was transferred into *E. Coli*. BL21 (DE3) strain. Cells were grown in 1 L LB media under 37 °C. IPTG (0.3 mM) was added to every 1 L medium in a shaker incubator to induce protein expression at 16 °C for 18 h. Cell pellets were harvested and lysed by French press with lysis buffer (50 mM Tris-HCl at pH 8.0, 250 mM NaCl, 0.5 mM TCEP), followed by centrifuging at 10000 rpm for 1 h. MgR protein was purified by passing through a column Nickle resin (GenScript Biotech Co.) and washed by using 3 column volumes of wash buffer (50 mM Tris-HCl at pH 8.0, 250 mM NaCl, 20 mM imidazole, 0.5 mM TCEP), then eluted with elution buffer (50 mM Tris-HCl at pH 8.0, 250 mM NaCl, 100 mM imidazole, 0.5 mM TCEP). The elution was concentrated and aliquoted into multiple small tubes and stored at −80 °C for further experiments. Protein was confirmed by SDS–PAGE gel, and its concentration was determined by standard BSA protein on the same gel.

### *In vitro* assay of RNA hydrolysis

Nm-modified ssRNA substrate was synthesized by GenScript Biotech Co. (5′6-Fam-UAACC UAUGA AGNNN UNmNNC UC-3′). Briefly, various purified proteins were reacted with 600 ng ssRNA substrate in reaction buffer (20 mM Tris-HCl pH 8.0, 100 mM KCl, 0.01 mM ZnCl_2_) at 37 °C for 30 mins. The reaction was terminated by heating to 85 °C for 10 mins. The products were separated by 20% urea-PAGE, visualized with a ChemiDoc XRS+\UnUniversal alHoodII gel imaging system (Bio-Rad) or Tanon 3500, and quantified with ImageJ software. A series of reactions with 400 ng ssRNA were designed to test the range of pH and reaction duration. Subsequently, multiple ssRNA substrates were incubated with 18 μg MgR in a 10 μl reaction (20 mM Tris-HCl pH 8.0, 100 mM KCl, 0.01 mM ZnCl_2_) at 37 °C for 30 mins to test MgR properties. The ssRNA sequences were as follows: ssRNA substrate with multiple modifications (5′6-Fam-AUACC GCAGC UAGAA UGAmGC Um^5^CAGA UGm^6^AAA-3′) and ssRNA substrate with hairpin structure (5′6-Fam-UAGCU CUGGA AAUUC UAGAmG CUAAU ACAAA-3′). Another Nm-modified ssRNA substrate was synthesized by GenScript Biotech Co. (5′-AGCCG CCUGG AUACC GCAGC UAGAA UGAmGC UGAGA UGAAA-3′) to test the consistency of hydrolysis. After reacting with MgR, reaction products were purified by an RNA Clean & Concentrator-5 kit (Zymo Research), followed by 5′ phosphorylation (New England BioLabs), repurification, and adaptor ligation (New England BioLabs). The final products were TA cloned and analyzed by Sanger sequencing. An Nm-modified ssRNA substrate was also synthesized by GenScript Biotech Co. (5′-AGCUA GNNNN NNNNN NNmNNN NNNNN NN-3′) and treated with the above protocol. The final products were high throughput sequencing.

### Cell culture, tissue and RNA extraction

HeLa cells, HEK293T cells, A549 cells, and Neuron-2a cells were purchased from the Shanghai Institute of Cell Biology, Chinese Academy of Sciences (Shanghai, China). All cells were grown in Dulbecco’s modified Eagle’s medium (DMEM) supplemented with 10% fetal bovine serum (FBS), 100 units/ml penicillin, and 100 μg/ml streptomycin at 37 °C with 5% CO_2_.

Following relevant guidelines and regulations, cancerous and distal noncancerous lung tissue samples were obtained from newly diagnosed lung cancer patients. Nanjing Drum Tower Hospital approved the study, and written consent was obtained from all patients.

Total RNA from cells and tissues was extracted by the Qiazol RNA extraction kit (QIAGEN). Extracted total RNA was confirmed by Nanodrop (Thermo Fisher) and 1% agarose gel electrophoresis (Sangon Biotech).

### Knockdown of FBL in HeLa cell line

For siRNA experiments, siRNA duplexes were used for fibrillarin silencing: 5′-GUCUU CAUUU GUCGA GGAA-AdTdT-3′ (sense sequence). Control siRNA does not target any human sequence (RiboBio). HeLa cells were transfected using LIPOFECTAMINE RNAIMAX reagent (Invitrogen) according to the manufacturer’s instructions and collected after 48 h. *FBL* mRNA expression was quantified by a HiScript II 1st Strand cDNA Synthesis Kit (+gDNA wiper) and ChamQ Universal SYBR qPCR Master Mix (Vazyme Biotech Co., Ltd.) according to instructions. The primers were listed in Appendix table 5.

### Viral infection

Neuro-2a cells were infected with MHV strain A59 at a multiplicity of 2 plaque-forming units (p.f.u.) per cell. The infected cells were cultured for 1 h and washed three times with PBS to remove the cell-free virus. The infected cells were harvested after 40h.

### rRNA elimination

Except for HeLa, HEK293T, and A549 cells, which used 15 μg total RNA, all the other samples were processed with 10 μg. The RNA was diluted to 29 μl, and 1 μl rRNA probe (H/M/R) was added (Vazyme Biotech Co., Ltd.). RNA was hybridized with the probe and heated to 95 °C for 2 min, then gradually cooled to 22 °C at 0.1 °C/s and maintained for 5 min. Next, the rRNA was eliminated by adding 1 μl RNase H (New England Biolabs), 4 μl 10X RNase H buffer, 4 μl DEPC water and incubating at 37 °C for 30 min. Subsequently, the hybridization and RNase H reaction were repeated 5/3 times to eliminate the rRNA fragment as much as possible. Finally, the probe was digested by 2 units of DNaseI (New England Biolabs) at 37 °C for 30 min. According to the manufacturer’s protocol, the RNA product was purified using VAHTS RNA Clean Beads (Vazyme Biotech Co., Ltd.).

### NJU-seq

RNA was fragmented by an RNA fragmentation reagents kit (Thermo Fisher) at 95 °C for 10 mins and then purified by 10 μl 3-M NaOAc (pH 5.2), 24 μl 5-μg/μl glycogen, and 372 μl 96% cold ethanol. For facilitating MgR binding of fragments with complex secondary structures after solubilization in water, every 1 μg RNA solution was ligated with 3 μl 25-μM poly(A)-ssRNA adaptor (pAGCUAAAAAAAAAAAAp, synthesized by GenScript Biotech Co.) at 16 °C overnight. The products were then reacted with 30 μg MgR in a 60 μl final volume with 20 mM Tris-Cl (pH 8.0), 100 mM KCl, and 0.01 mM ZnCl_2_ at 37 °C for 30 min. Next, the products were purified by an RNA Clean & Concentration-5 kit (Zymo Research), followed by 5′ phosphorylation (39 μl RNA, 5 μl (10X T4 PNK Reaction Buffer), 5 μl ATP (10 mM), 1 μl T4 PNK (3′ phosphatase minus, New England BioLabs)). Finally, we constructed the NGS library using the products from the last step following the NEBNext Small RNA Library Prep Set for Illumina (New England BioLabs) protocol. Fragments smaller than 50 nt were extracted from a 6% PAGE gel and sent for NGS using Illumina X-Ten. The exact process prepared the NC sample (negative control group) without MgR protein.

### NGS data analysis and statistics

Nm site detection and various statistical analyses were performed using homemade Perl and R scripts and open-source bioinformatics tools. Adaptors were filtered from raw reads using Cutadapt (Version 2.10). Reads were mapped the reference sequences by Bowtie 2 (Langmead & Salzberg, 2012) (Version 2.3.4). rRNA reference sequences are shown in Appendix table 1, and the transcript reference sequences were from GENCODE release 34 and M25 (Frankish et al., 2019). Only protein-coding transcripts with ‘BASIC’ tags and without ‘cds_start_NF’ or ‘cds_end_NF’ tags were retained. Perl scripts handled alignment results, including filtering low-quality or multiple-hit alignments and calculating each site’s reads’ end count. The likelihood of 2′-O-methylation modification at each site was judged based on the following scoring system.

### Scoring 2′-O-methylation modification

As shown in Appendix figure 5, we counted the stacking position of the 3′ end of the reads for the treatment group with MgR digestion (TR) and the control group without digestion (NC). First, normalization correction was performed according to the total aligned reads. Then, scoring was performed for each site on the rRNA and transcriptome reference based on the number of reads ending at the current position (N) and the latter (N+1). The NJU-seq score algorithm was as follows.

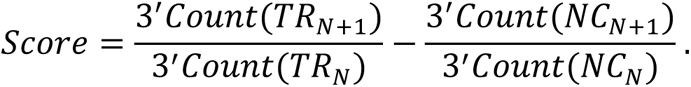

The average of the three replicates’ scores was taken and ranked to complete identification for modifications, and here, we took 30 as a threshold. To prevent some points with a similar high N+1 enrichment in NC from being misclassified, we also required 3’*Count*(*TR_N_*_+1_) > 3 × *mean*(3’*Count*(*NC_N_*_+1_)). At this point, we finished processing the NJU-seq data and acquiring the Nm sites for subsequent analysis.

### Statistics on genomic features

We mainly used Perl and R scripts to perform tri-ribonucleotide unit statistics, presentation of distribution positions, and alternative splicing region statistics on all Nm sites.

Sequence probability logo plots for Nm sites were drawn by the R package *ggseqlogo* (Wagih, 2017). The length-normalized mRNA transcript was obtained by averaging all Nm-located transcripts’ 5′UTR/CDS/3′UTR length of each sample. Equal length bins of three regions were taken for Nm site counting in each standard transcript. We counted the distance from the Nm site position to the start and stop codon. ASE and CSE were obtained from the statistics of all alternative splicing events provided in the SpliceSeq database (Ryan et al., 2012).

### Quantitation of RNA Nm status by RTL-P method

Two sets of primers were designed for the target site, with the F_U_ forward primer upstream of the Nm site and the F_D_ forward primer downstream of the Nm site (Appendix table 5). For rRNA Nm detection, the high dNTP concentration reaction mixture consisted of 5× RT Buffer 4 μl, M-MLV (H-) Reverse Transcriptase (Vazyme Biotech Co., Ltd., 200 U/μL) 1 μl, RNase inhibitor (40 U/μL) 1 μl, RNA 100 ng, RT primer (10 μM) 1 μl, RNase-free H_2_O 5 μl, dNTPs (1 mM each) 1 μl. The low dNTP concentration reaction mixture was replaced by dNTPs (2 μM each). Then, the mixtures were incubated at 45 °C for 1 hour and 85 °C for 2 min. The cDNA was diluted 100-fold and subsequently subjected to qPCR, with both F_U_ and F_D_ products amplified for each cDNA. For mRNA Nm detection, 1 μg of total RNA was used for further RTL-P tests. RT efficiency and RT fold change were calculated according to the following strategy. RT efficiency=template amount measured by F_U_ and R/template amount measured by F_D_ and R. RT fold change=RT efficiency with low dNTPs/RT efficiency with high dNTPs (Dong et al., 2012).

### The assay of RNase H cleavage

GenScript Biotech Co synthesized RNA oligonucleotides and chimera probes. Briefly, 12.5 pmol 5**′**-FAM-labeled RNA oligonucleotides were mixed with a 75 pmol chimera probe, heated to 95 °C for 2 min, cooled to 22 °C at 0.1 °C/s, and maintained for 5 min. Next, the hybrid was reacted with 1 μl RNase H (New England BioLabs) at 37 °C for 30 min and then heated to 90 °C for 10 min to terminate the reaction. The cleavage products were added to RNA Dye (New England BioLabs), analyzed by 20% UERA-PAGE, and visualized by a ChemiDoc XRS+\UnUniversal alHoodII gel imaging system (Bio-Rad).

### Nm-VAQ assay of RNA 2′-O-methylation ratio

Synthetic RNA oligonucleotides with/without Nm were obtained from GenScript Biotech Co., and Sangon Biotech Co. synthesized qRT–PCR primers. All sequences were listed in Appendix table 5. The oligonucleotides with/without Nm were mixed to obtain gradient 2′-O-methylation ratio substrates. Then, 6.25*10^-2^ pmol substrate was added to 6.25 pmol D(4)R(13) chimera probe at 95 °C for 2 min, followed by cooling to 22 °C at 0.1 °C/s and 22 °C for 5 min. The products were divided into 2 parts for the following reaction. One mixture contained 5 μl previous product, 1 μl RNase H (New England BioLabs), 1 μl 10X RNase H Reaction Buffer, and 3 μl RNase-free H_2_O. RNase H storage buffer was substituted for RNase H to form the other mixture to serve as a blank control, followed by 30 min at 37 °C and 10 min at 90 °C. The products were diluted 50-fold and then used to obtain cDNA by a HiScript II 1^st^ Strand cDNA Synthesis Kit (Vazyme Biotech Co., Ltd.). cDNA was diluted 100-fold, and then qPCR was conducted in a 20 μl reaction mixture containing 10 μl ChamQ Universal SYBR qPCR Master Mix (Vazyme Biotech Co., Ltd.), 0.4 μl F primer, 0.4 μl R primer, 2 μl cDNA, and 7.2 μl H_2_O, following the protocol: 30 s at 95 °C, then 40 cycles of 95 °C for 10 s and 60 °C for 30 s. Each cDNA was analyzed in 3 replicates. ΔCT (cycle threshold) = Ct value of RNase H reaction-Ct value of control. The linear relationship between the Nm ratio and ΔCT was obtained by GrapicPrism. The additional 6.25 pmol, 0.625 pmol, and 0.0625 pmol substrates with 50% Nm ratio were also assayed (cDNA did not dilute), and ΔCT was obtained. The measured modification ratios were calculated by the above linear relationship.

### Quantitation of RNA Nm status by Nm-VAQ method

All primers were obtained from Sangon Biotech Co., and the sequences are shown in Appendix table 6. For rRNA Nm detection, 100 ng total RNA and 6.25 pmol chimera probes (molar ratio of rRNA molecules to chimera probes was ∼1:200) were hybridized, followed by RNase H cleavage, reverse transcription, and qPCR (cDNA was diluted to 100-fold) according to Nm-VAQ protocol. ΔCT was used to calculate the ratio of modification. In addition, some treatments were adjusted to perform mRNA Nm detection. 2-10 μg of total RNA was used for hybrids, and 4 μl of cDNA was used for the qPCR assay.

### *In vitro* transcription

The Nm-VAQ detection DNA fragment of *PLXNB2* (22, 50,290,219, Gm) and *BCL6* (3, 187,721,452, Gm) were obtained by qRT-PCR. The T7 primer (TAATA CGACT CACTA TAGGG) was then added to the DNA fragment by PCR. The DNA fragments were as templates to produce RNA fragments without modifications following the T7 High Yield RNA Transcription kit instructions (Vazyme Biotech Co., Ltd.). 1ng of *in vitro* transcribed RNA was mixed with 1μg total Arabidopsis RNA, followed by hybridization, RNase H cleavage and reverse transcription according to the Nm-VAQ. The cDNA was amplified by PCR and then the product was detected by electrophoresis. The *in vitro* transcriptome total HeLa mRNA was generated as described in the Zhang et al., study (Zhang et al., 2021). The modification-free mRNAs were then performed NJU-seq process and calculated the score.

### The snoRNA target prediction

The C/D box snoRNAs were downloaded from snoPY (http://snoopy.med.miyazaki-u.ac.jp). All 427 C/D box snoRNA were obtained. The efficient target prediction for box C/D snoRNAs was followed by PLEXY (http://www.bioinf.uni-leipzig.de/Software/PLEXY/).

### qRT–PCR and calculation of expression of NCAM1 transcripts

1 μg of total RNA was used to obtain cDNA with a HiScript II 1st Strand cDNA Synthesis Kit (Vazyme Biotech Co., Ltd.). cDNA was added to ChamQ Universal SYBR qPCR Master Mix (Vazyme Biotech Co., Ltd.), following the process: 95 °C for 30 s, 40 cycles of 95 °C for 10S and 60 °C for 30 s. Each cDNA was assayed in 2 replicates. The primer oligos were as follows: (NCAM1-all-F: CAACC TTGGG AGGCA ATTCT), (NCAM1-all-R: ACTGC CATTA AAAAG GGGGC), (NCAM1-others-F: ACCTT GGGAG GCAAT TCTGC), and (NCAM1-others-R: GCAGA AACTT CTCTG TAAAT CTAGC). The standard curve was completed as above. Standards of qRT–PCR amplification products were subjected to gradient dilution. qPCR was completed under standard protocols based on the above reagents and procedures.

### RNA-seq

We depleted rRNA from total RNA, fragmented it into ∼300 bp fragments, and prepared the library for RNA-seq. After library preparation and pooling of different samples, the samples were subjected to PE150 sequencing on the Illumina NovaSeq 6000 platform. Paired-end reads were aligned to the GENCODE M25 genome using Bowtie2. Gene expression statistics were performed using HTSeq (Anders et al., 2015) (Version 1.99.2), and differentially expressed genes were obtained using DESeq2 (Love et al., 2014) (Version 1.30.1).

### Protein abundance

The HeLa protein abundance data were downloaded from the Protein Abundances Across Organisms (PaxDb, Experiment ID 182) (Geiger et al., 2012; Wang et al., 2015). We used transcriptome expression data from the ENCODE (Davis et al., 2018) portal with the following identifiers: ENCFF796REI and ENCFF846THO, and only kept genes with TPM > 0.2. The 6,881 genes with both protein abundance and mRNA expression were used for subsequent analysis. Among them, the number of Nm-located genes was 1,397.

### Gene ontology (GO) enrichment

For functional enrichment analysis, genes containing Nm sites were uploaded to DAVID Bioinformatics Resources (http://david.abcc.ncifcrf.gov). Enriched GO terms were restricted to P ≤ 0.05.

### Simulation of structural models

The influence of 2′-O-methylation on RNA-RISC binding was simulated by PyMOL (Version 2.4). We used the molecular structure of Argonaute2-miR-122 bound to the target RNA (PDB ID: 6MDZ) (Sheu-Gruttadauria et al., 2019) for observation and further simulated the modification at some target RNA sites. The methyl group bond angle was strictly controlled to match the real situation, and the distance to the surrounding amino acids before and after binding was measured.

## Data and code availability

All data are available in the National Center for Biotechnology Information (accession nos. SAMN17082771 to SAMN 17082794 and SAMN17121988 to SAMN17122043). In addition, principal analysis codes are available on https://github.com/IvanWoo22/NJU_seq.

## Acknowledgments

We are grateful to the High-Performance Computing Center of Nanjing University for performing the numerical calculations on its blade cluster system in this paper. In addition, we acknowledge all members of Dr. Chen Jian-Qun’s lab, Dr. Li Shanqing and Dr. Liang Naixin for comments and discussions. We also gratefully acknowledge Dr. Chan Robin, Dr. Zhang Hui, and Dr. Zhou Li for editing the paper.

## Funding

This work was supported by the Youth Program of the National Natural Science Foundation of China (31801065), the National Key R&D Program of China (2021YFC2701603), Fundamental Research Funds for the Central Universities (021414380507), and National Natural Science Foundation of China (32170218).

## Author contributions

Q.H.C., Q.W., and J.Q.C., designed this research; Y.T., S.N.W., X.L.L., X.W.G., Y.L., F.Y., R.L.X., Y.W., and L.W.L. performed this research; Q.H.C., Q.W., Y.T., and Y.F.W., wrote the manuscript; Y.F.W. and Q.W., analyzed the data; T.W., and Z.C.J., provided the clinical samples and pretreatment.

## Competing interests

No competing interests declared.

## Notes

### Competing Interest Statement

The authors have declared no competing interest.

### Summary of Updates

The proof of the authenticity of the identification site is reliable evidence for the detection tool. In the new version, we added more evidence to detect the Nm status of mRNA sites by NJU-seq. The following results were obtained: 1) 4074 Nm sites were identified in HeLa cell line by NJU-seq with restrict standard, in which 52 of 67 selected sites were further validated to be 2′-O-methylated more than 1%. Unlike rRNA, the 2′-O-methylation ratios of most validated Nm sites on mRNA from cell lines were lower than 30%, which was out of the detection abilities of previous methods. Although some screened sites were proved to be methylated with lower than 1%, the quantity of methylated fragments was not negligible considering their high expression level. 2) In vitro transcriptome mRNA was used as a negative control to confirm the results were not caused by secondary structures. 3) Based on the previous studies that RNase H was inhibited by Nm, Nm-VAQ was developed for further validation. 4) Multiple in vitro assays and controls in each experiment. We believe updating these new results can better help readers.

