## Supplemental Figures for "A novel platform of RNA 2′-O-methylation high-throughput and site-specific quantification tools revealed its broad distribution on mRNA"

### Appendix information

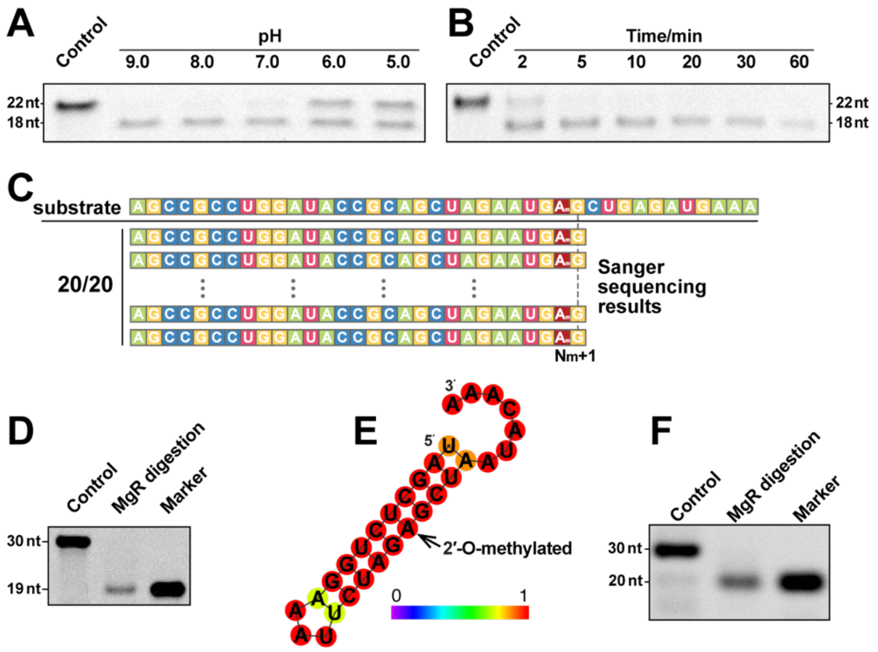


Appendix figure 1 Recognition of MgR and Specificity of NJU-seq for 2′-O-methylation modifications.

(A and B) Exploration of MgR hydrolysis conditions *in vitro*. The ssRNA substrate (5′-6-Fam-UAACC UAUGA AGNNN UNmNNC UC-3′, N represents the mix of A/G/C/U) was reacted with a series of different conditions as shown. A, pH ranges from 5.0 to 9.0; B, Reaction time ranges from 0-60 min.

(C) TA clone MgR hydrolyzed ssRNA products. The ssRNA substrate (5′-AGCCGCCUGG AUACCGCAGC UAGAAUGAmGC UGAGAUGAAA-3′) was reacted with MgR protein, followed by TA cloning into *E. coli*. Twenty insert sequences from independent clones were sequenced by Sanger sequencing. All products ended 1 nt downstream of the Am site.

(D) Recognition specificity of MgR for Nm modifications. MgR digested the ssRNA substrate (5′-6-Fam-AUACC GCAGC UAGAA UGAmGC Um$^5$CAGA UGm$^6$AAA). The marker band responded to the Nm+1 site.

(E) Testing of MgR to hydrolyze complex RNA structure. RNAFold predicted the substrate (5′-6-Fam-UAGCU CUGGA AAUUC UAGAmG CUAAU ACAAA-3′) structure.

(F) MgR-digestion products of the substrate in Figure S1E.

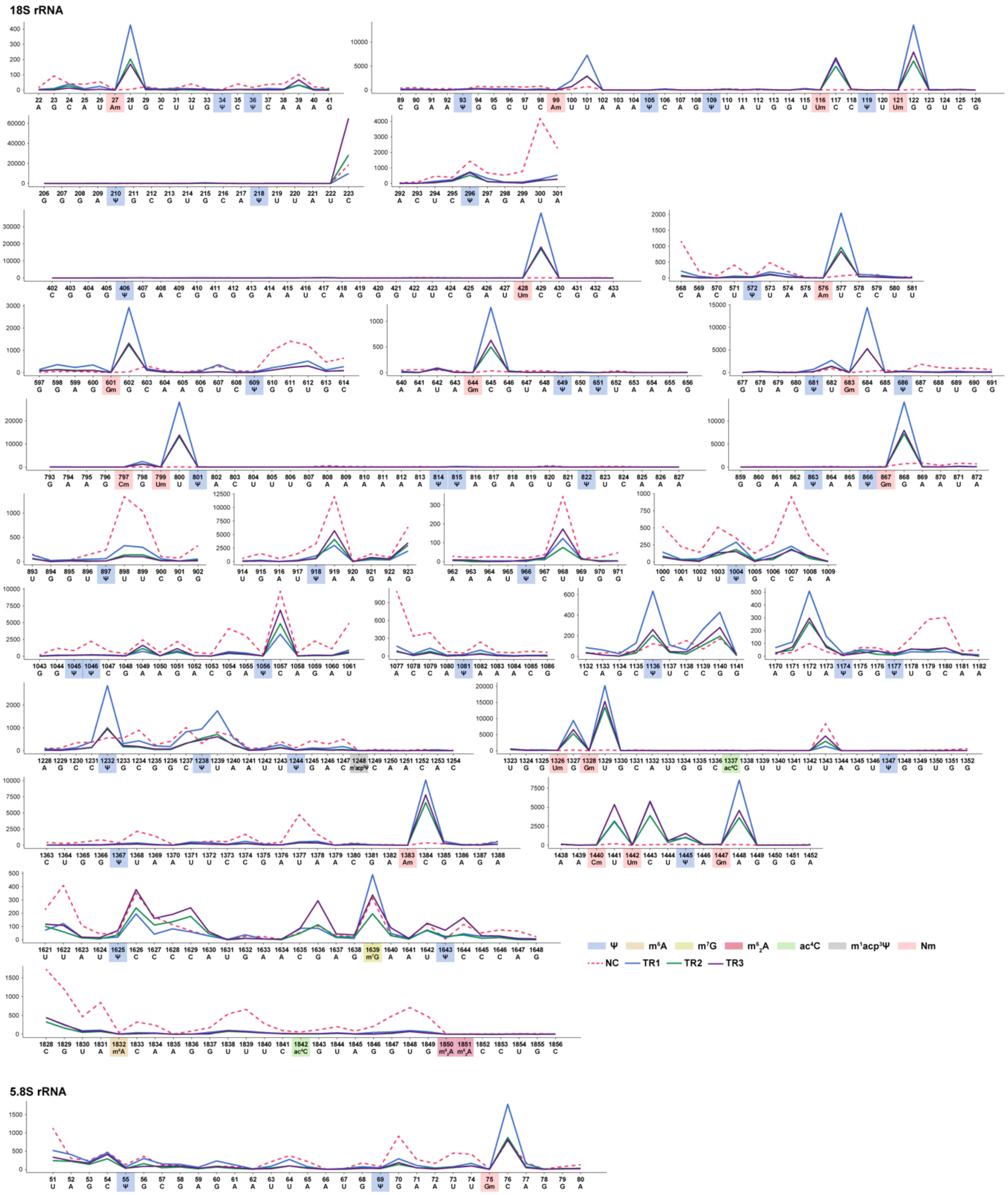


Appendix figure 2 3′ end counts on other modifications of HeLa 5.8S and 18S rRNA with (TR, MgR treatment, solid line) and without (NC, dotted line) MgR digestion. The different colors marked the presence of different RNA modifications in previous reports.

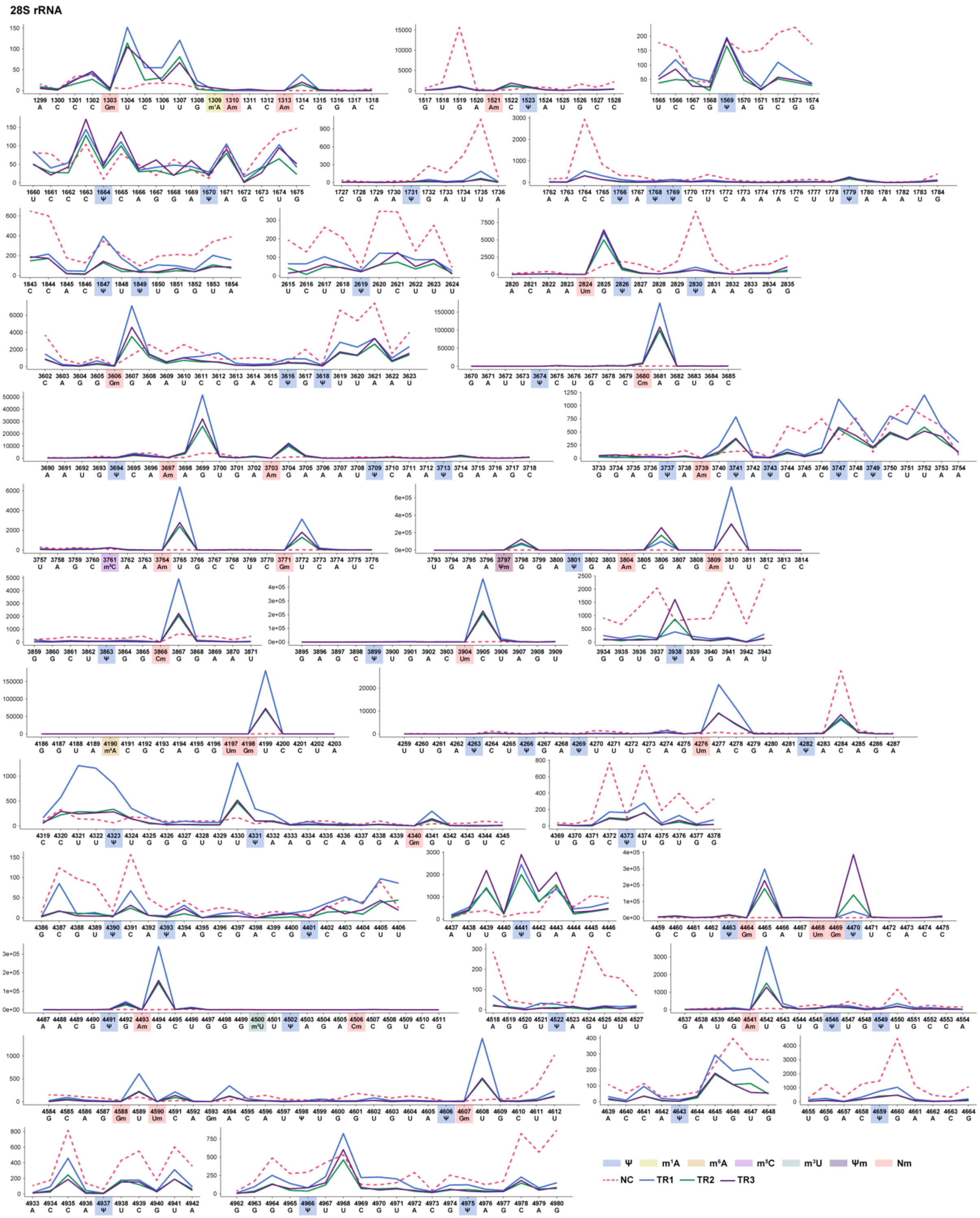


Appendix figure 3 Other kinds of RNA modifications on HeLa 28S rRNA.

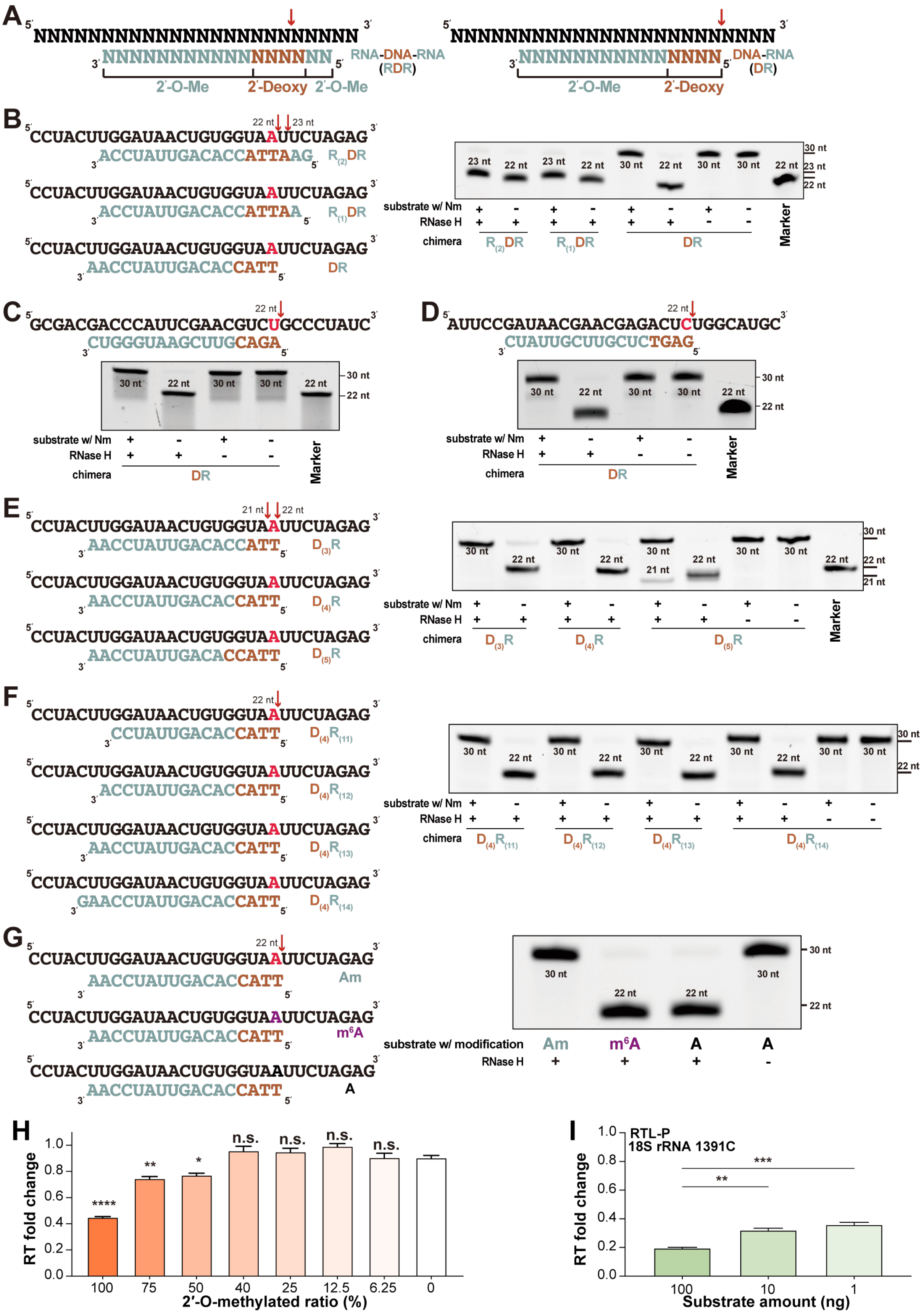


Appendix figure 4 The construction of Nm-VAQ, and RTL-P detection of Nm sites with known methylated ratio. Screening chimera probe for RNase H-dependent Nm detection.

(A) Schematic presentation of chimera probe. The red arrow indicated the RNase H cleavage sites in a previous report. The baby blue sites indicated the RNA of the chimera probe, which all were 2′-O-methylated, and the reddish-brown sites indicated the DNA of the chimera. The following chimeras were labeled in the same way.

(B-F) The exploration of chimera probe structure. The scheme of hybrid of RNA substrate (up sequences. All were labeled with FAM at the 5′ end), and chimera probes (down sequences) was shown on the left. The RNase H reaction products were presented on the right by electrophoresis. The red sites in the substrates indicated the Nm site, and the red arrows indicated the cleavage sites. The number presented the length of FAM-labeled cleavage products.

(B) Site-specific cleavage of RNase H directed by RDR or DR chimeras. Although both R(1)DR (with one ribonucleotide at 5′ end) and R(2)DR (with two ribonucleotides at 5′ end) chimera probes induced specific cleavage sites on the unmethylated substrate and produced 22 nt products, they were not inhibited by 2′-O-methylation completely. On the other hand, a chimera probe designed with only DR (without ribonucleotides at 5′ end) demonstrated clear site-specific cleavage on unmethylated substrates but not 2′-O-methylation substrates.

(C and D) Cleavage of RNase H in the other two substrates. The DR chimera probe also inhibited the RNase H cleavage activity.

(E) RNase H cleavage directed by DR chimera with varied DNAs. The cleavage activity of probes D(3)R and D(4)R was wholly inhibited by 2′-O-methylation. Due to the higher stability of DNA over RNA, D(4)R was chosen for subsequent testing.

(F) RNase H cleavage directed by DR chimera with varied RNA numbers. There was no significant difference in cleavage sites or cleavage efficiency among tests with different chimera probes. In principle, the longer length of RNA required a higher melting temperature, facilitating the test of Nm sites in strong secondary structural regions.

(G) Effect of $m^6A$ and Nm modification on RNase H cleavage. The purple sites were $m^6A$-modified. The $m^6A$-containing substrate was cleaved by RNase H at the modification site, while Am inhibited the cleavage completely.

(H) $6.25*10^{-2}$ pmol synthesized substrate with and without Nm were mixed to obtain a known Nm ratio. See Methods for calculating RT fold change, negative with Nm ratio. All results were compared with a 0% Nm ratio (negative control). *$P \leq 0.05$, **$P \leq 0.01$, ****$P \leq 0.0001$, n.s., not significant, one-way ANOVA.

(I) The RT fold change of 18S 1391Cm site among three total RNA amounts detected by RTL-P. **$P \leq 0.01$, ***$P \leq 0.001$ by one-way ANOVA. Error bars describe SEM for three technical replicates.

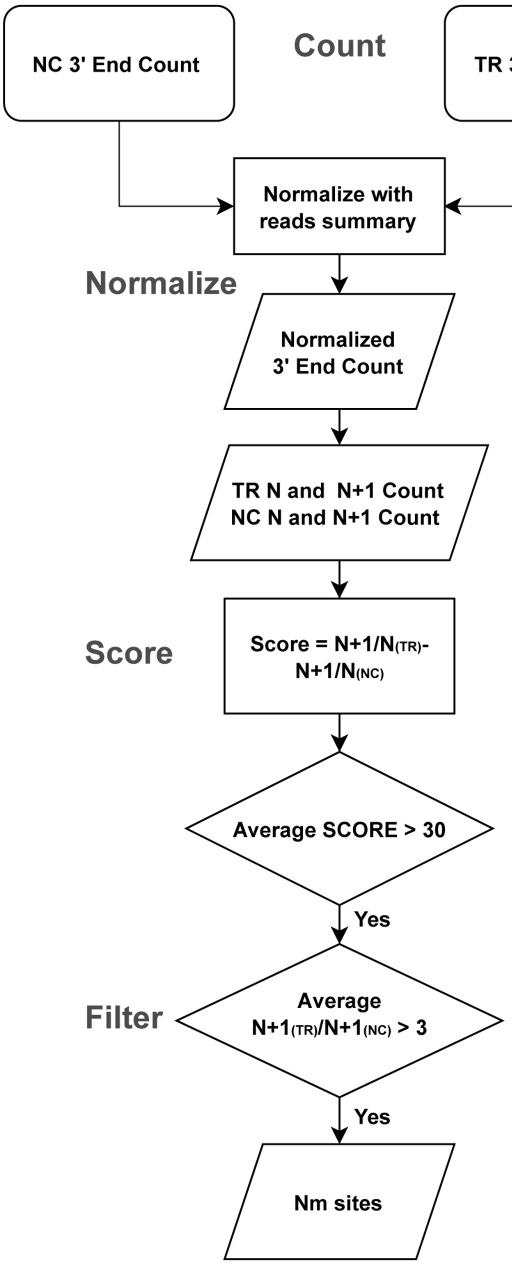


Appendix figure 5 Scoring and filtering process for NJU-seq.

According to this flow, Nm sites would be detected from the reads aligned to the reference sequence.

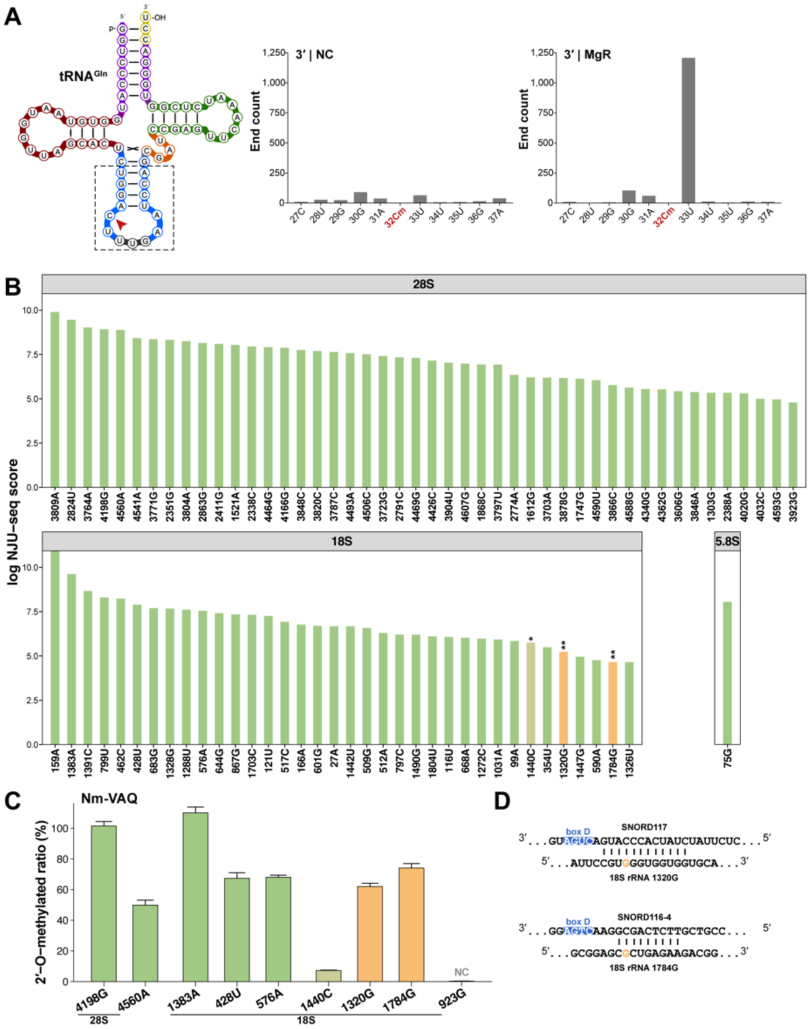


Appendix figure 6 Identification of transfer RNA (tRNA) and Ribosomal RNA (rRNA) Nm sites using NJU-seq and Nm-VAQ.

(A) Plot of reads 3′ end counts at 32Cm of tRNA-Gln-UUG without and with MgR digestion.

(B) The Nm sites identified successfully on HeLa rRNA by NJU-seq. Nm sites in green were identified in previous multiple studies, and sites in orange were newly identified in 2020.

(C) Selected site validation by Nm-VAQ. Y-axis represented the 2′-O-methylation ratios of each site. Non-modification-reported 18S 923G served as a negative control.

(D) Potential snoRNA targeting of the 18S 1320G and 1784G sites.

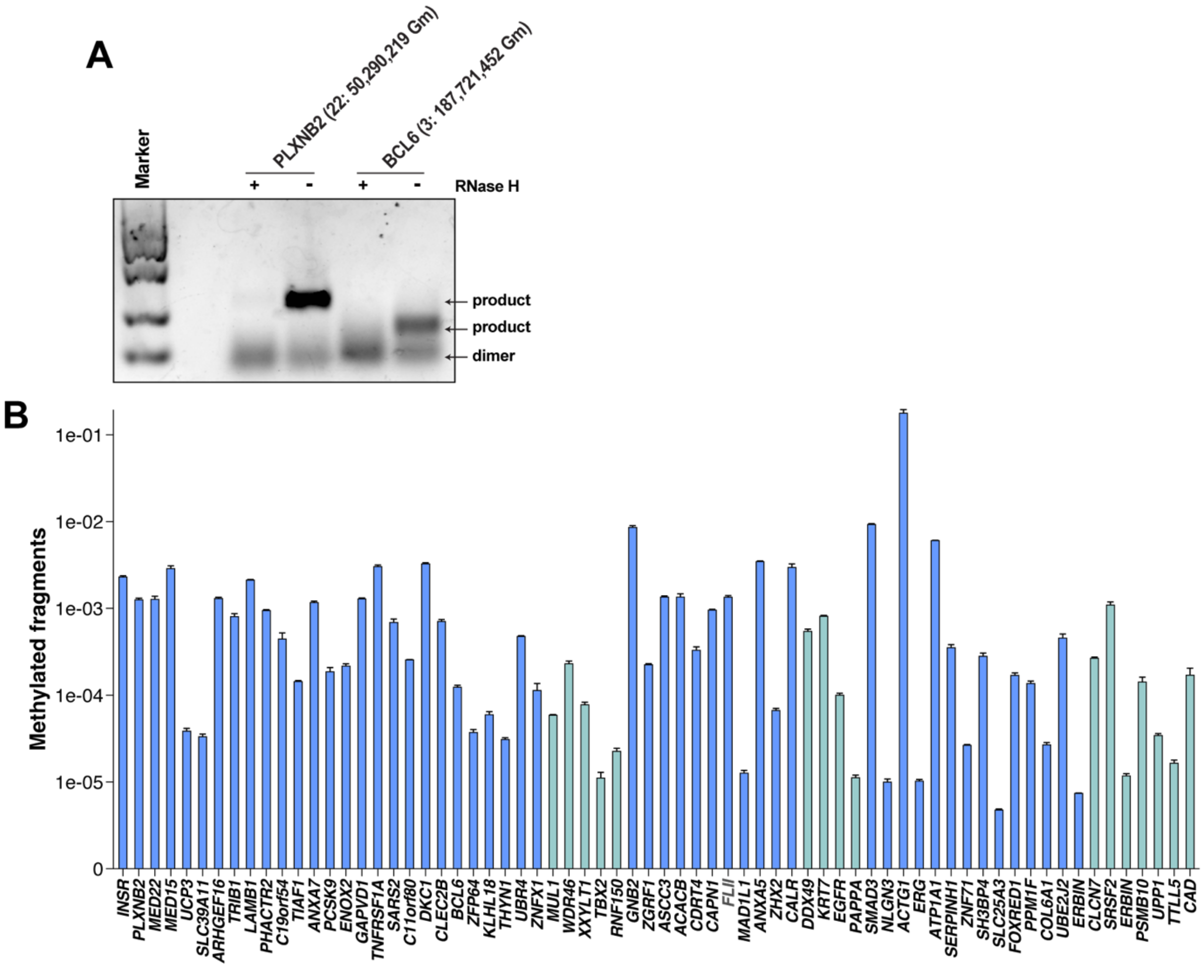


Appendix figure 7 Nm-VAQ detection of transcribed RNA fragments *in vitro* and the methylated RNA fragments by Nm-VAQ.

(A) The Nm-VAQ detection DNA fragment of *PLXNB2* (22, 50290219, Gm) and *BCL6* (3, 187721452, Gm) as templates for *in vitro* transcription. The unmodified RNA fragments were obtained, followed by hybridization with the probe, RNase H cleavage, and reverse transcription. Finally, the cDNA was obtained for PCR amplification, and the products were detected by electrophoresis.

(B) The gene mRNA expressions were calculated by the $2^{-\Delta CT}$ method and normalized to *GAPDH*. Subsequently, the number of methylated fragments was obtained by multiplying the proportion of methylation with the expression.

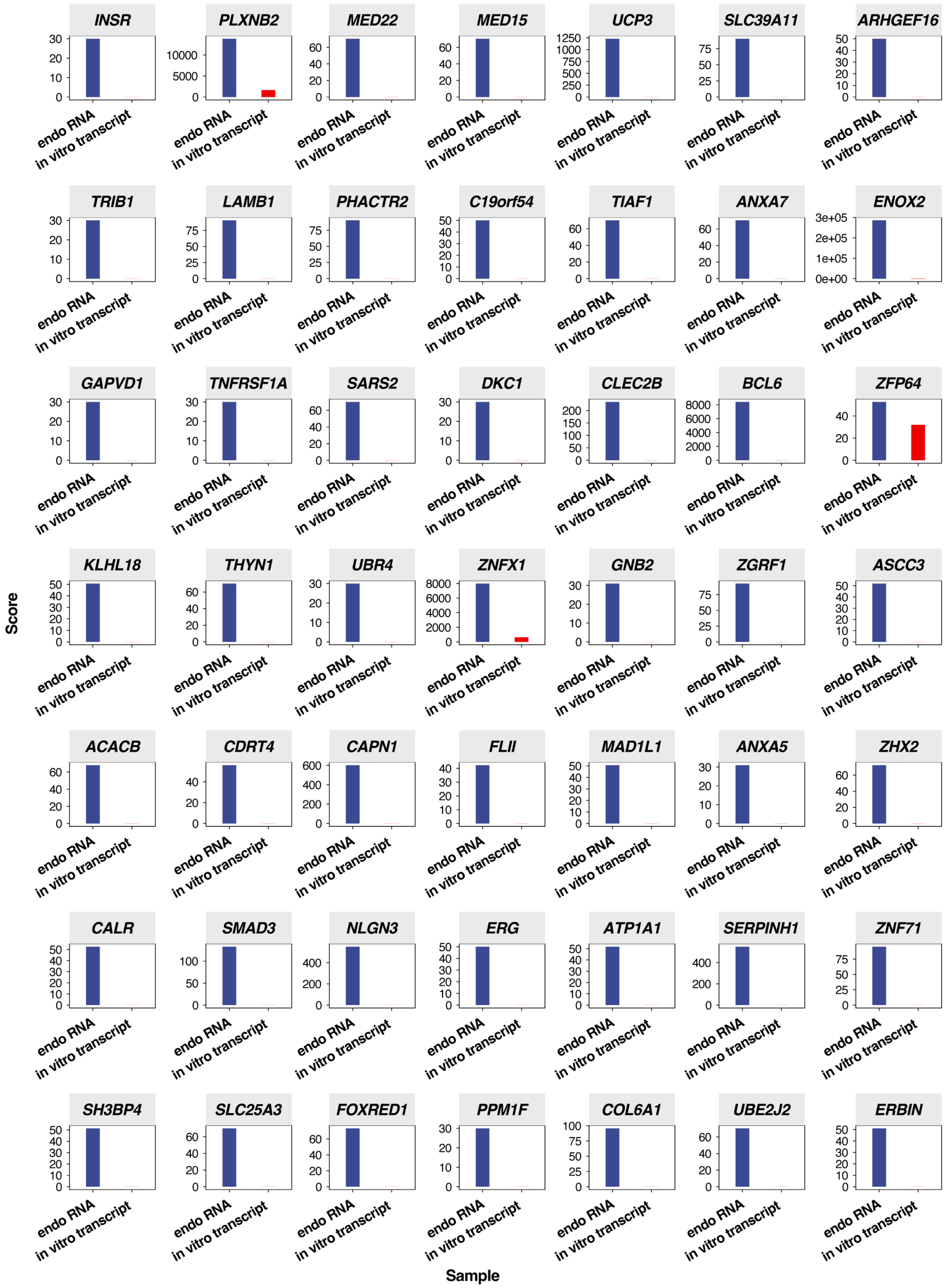


Appendix figure 8 The mRNA site scores of *in vitro* transcriptome. mRNA Nm sites score of endo HeLa mRNA (left), and *in vitro* transcriptome (right).

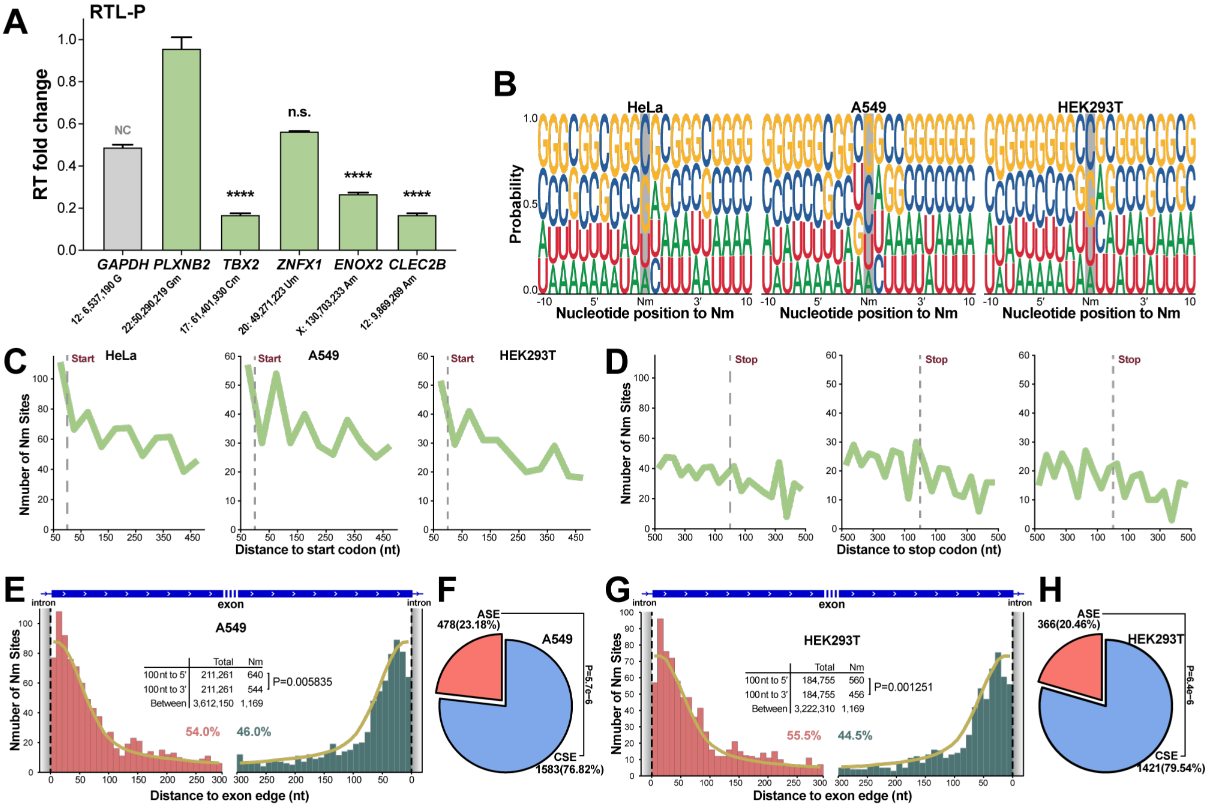


Appendix figure 9 Selected HeLa mRNA Nm status detected by RTL-P, and mRNA Nm site distribution in HeLa, A549, and HEK293T cells.

(A) *GAPDH* (12:6,537,190, G) was used as a negative control. Error bars describe SEM for three technical replicates.

(B) Sequence probability logo for mRNA Nm sites. Nm sites were marked in gray background.

(C and D) Distribution of three cell lines' Nm sites in windows with absolute length centered on the start (C) and stop (D) codon.

(E) Distribution of A549 Nm sites near exon 5′ (left) and exon 3′ (right) boundaries.

(F) Distribution of A549 Nm sites in ASE and CSE regions was shown.

(G) Distribution of HEK293T Nm sites near exon 5′ (left) and exon 3′ (right) boundaries.

(H) Distribution of HEK293T Nm sites in ASE and CSE regions was shown.

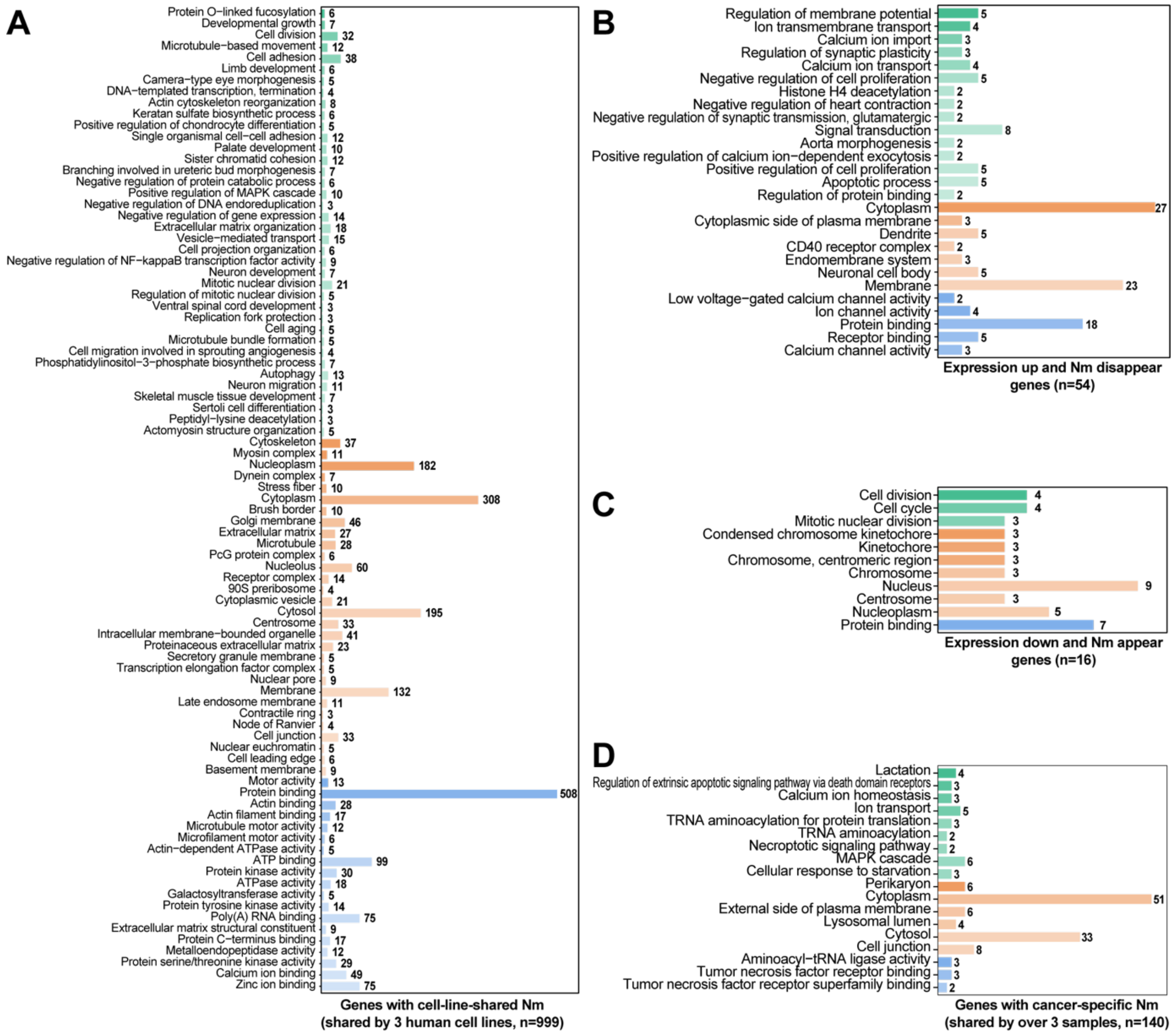


Appendix figure 10 GO analysis of Nm-modified genes.

(A) GO analysis of genes containing Nm sites shared by HeLa, A549, and HEK293T.

(B) GO analysis of up-regulated genes that contained Nm sites disappeared after MHV infection.

(C) GO analysis of down-regulated genes which contained Nm sites appeared after MHV infection.

(D) GO analysis of genes with cancer-specific Nm sites. Only the Nm site shared by over three cancer samples was analyzed.

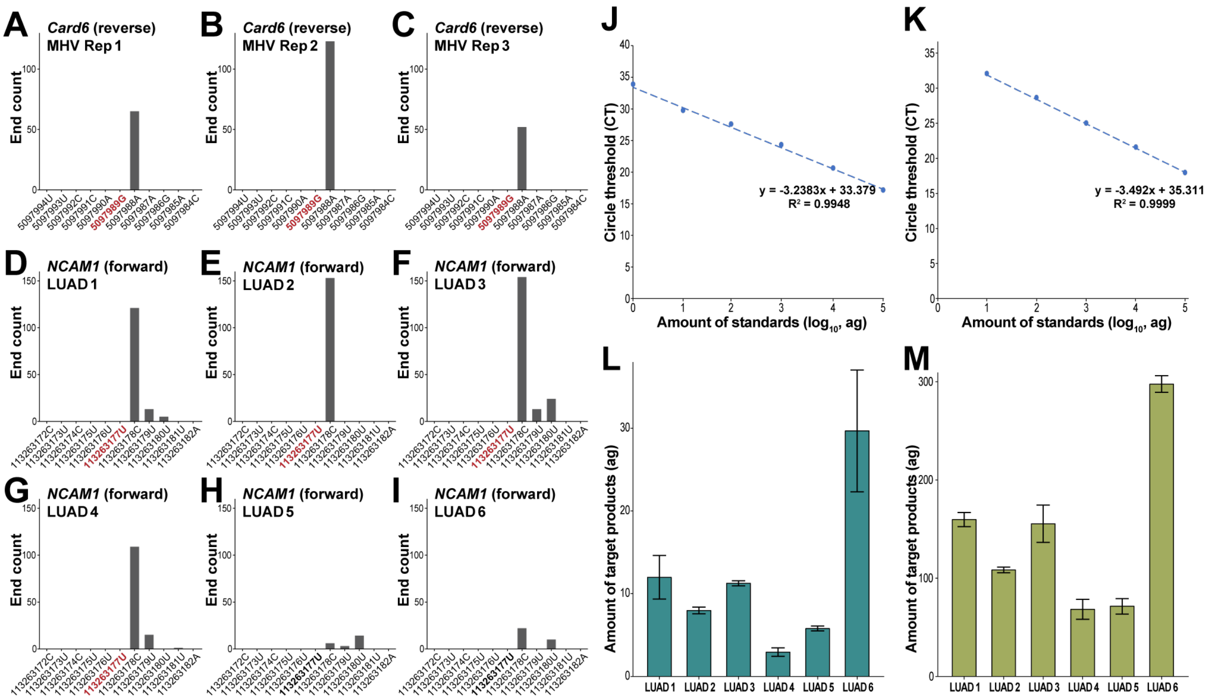


Appendix figure 11 End count on the Gm site of *Card6* and the Um site of *NCAM1*, and expression of *NCAM1* transcripts covering the Nm site.

(A-C) Plot showing reads' 3′ end count between positions 15:5,097,984 to 15:5,097,994 of the mouse genome.

(D-I) Plot showing reads' 3′ end count between positions 11:113,263,172 to 11:113,263,182 of the human genome. Sites marked with red were detected as Nm sites.

(J) Standard curve for amounts of transcripts other than NCAM1-211 standards and cycle threshold (CT) values of qPCR.

(K) Expression of transcripts other than NCAM1-211.

(L) Standard curve for amounts of all transcripts containing the Um site standards and CT values of qPCR.

(M) Expression of all transcripts. The calculation process was detailed in Methods. Error bars describe SEM for two technical replicates.

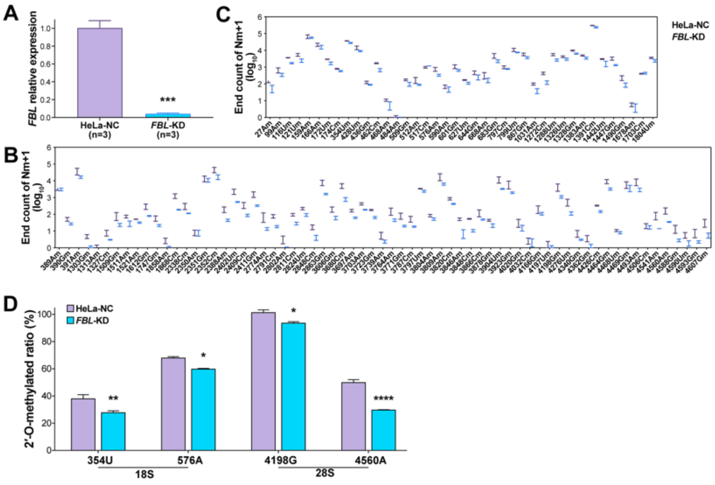


Appendix figure 12 NJU-seq is sensitive to Nm modification.

(A) Expression of *FBL* mRNA. *GAPDH* was used as an internal reference gene. *FBL*-KD mRNA expression was calculated by normalization to the HeLa-NC group. Error bars describe SEM (n = 3). ***P<0.001 (unpaired t-test).

(B and C) The read end count of sites 1 nt downstream previously reported Nm in 18S (B) and 28S rRNA (C) of HeLa-NC and *FBL*-KD samples by NJU-seq (n = 3). The end count was normalized according to total aligned reads.

(D)The methylation ratio of four sites previously reported as Nm sites in wild type and *FBL*-KD group HeLa cell line by Nm-VAQ. Y-axis represented the 2′-O-methylation ratios of each site. *P<0.05, **P<0.01, ****P<0.0001, by unpaired t-test. Error bars in 2C-2E described SEM for three technical replicates.

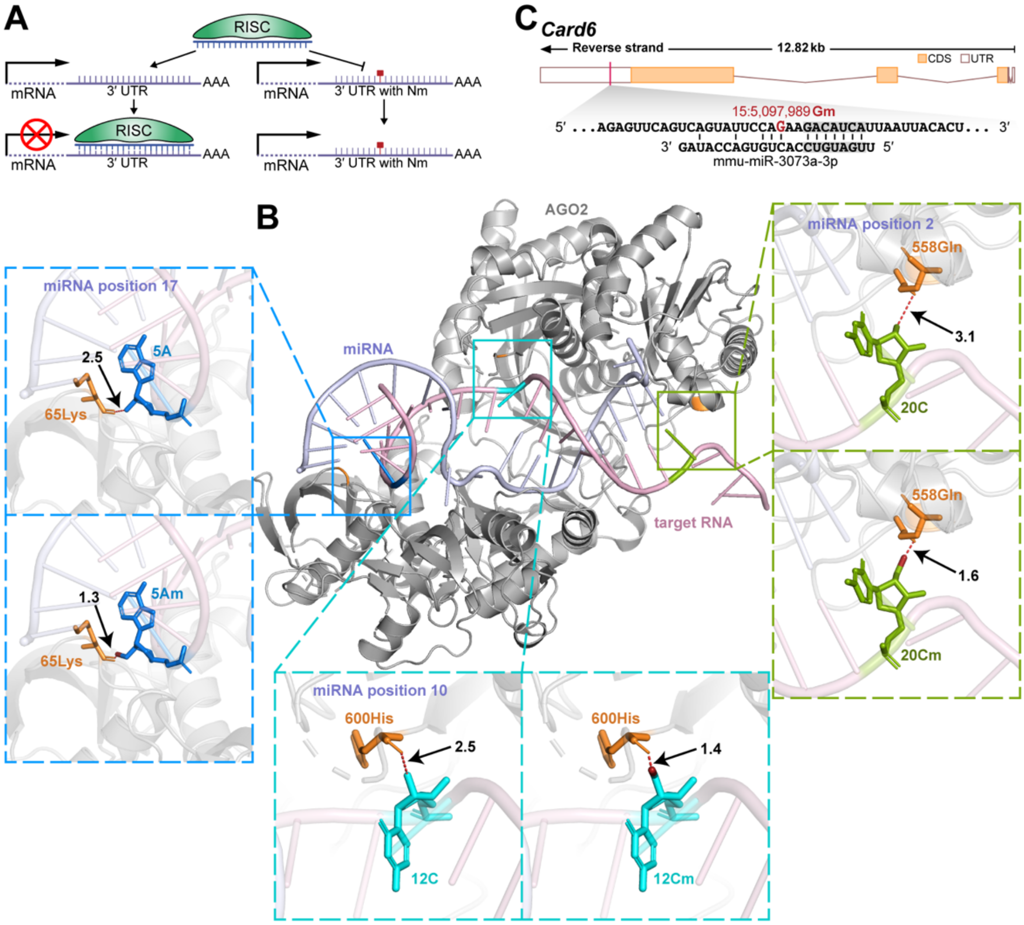


Appendix figure 13 Nm sites might affect the binding and cleavage of RISC to target RNA.

(A) Molecular model of Nm site influencing RISC binding mRNA.

(B) Schematic representation of the binding of the human Ago2-miR122 (blue) RISC complex to target RNA (pink) (based on PDB ID 6MDZ). Target RNA's 5A, 12C, and 20C ribose 2-hydroxy bind to Ago2 65Lys, 600His, and 558Gln via hydrogen bonding, corresponding to positions 17, 10, and 2 of miRNA, respectively. Nm might affect the binding of RISC and target RNA. The shortening of the distance between ribulose-2′-O-$CH_3$ and the corresponding amino acid of Ago2 affects hydrogen bond formation after simulating the occurrence of a 2′-O-methylation modification at the target RNA site.

(C) Location of a Gm site, 15:5,097,989, on *Card6* mRNA might prevent binding with miR-3073a-3p. The seeding region was marked in gray.

Appendix table 1 NJU-seq identification of rRNA Nm sites in HeLa cell lines.

Appendix table 2 NJU-seq identification of mRNA Nm sites in 3 human cell lines.

Appendix table 3 NJU-seq identification of mRNA Nm sites in normal and MHV- infected Neuro-2a cell lines.

Appendix table 4 NJU-seq identification of mRNA Nm sites in 8 pairs of cancer samples.

Appendix table 5 Primer sequences and chimera probe sequences used in this study.

Appendix table 6 The methylation ratio of selected Nm sites by Nm-VAQ detections.
